# CD11c+ Tbet+ B cells constrain obesity- and vaccination-induced germinal center B cells and T helper cells

**DOI:** 10.1101/2025.09.01.673552

**Authors:** Carlo Vanz, Benjamin T. Enslow, Emma Collins, Madilyn Dominguez-Lowry, Nathaniel Liendo, Elizabeth A. Dudley, Elizabeth A. Leadbetter

**Affiliations:** Department of Microbiology, Immunology & Molecular Genetics, UT Health San Antonio; San Antonio, Texas, USA; Department of Health Sciences, UT Health San Antonio; San Antonio, Texas, USA; University of Texas San Antonio; San Antonio, Texas USA; St. Mary’s University, San Antonio, Texas, USA

**Keywords:** CD11c+Tbet+ B cells, obesity, immune complexes, T_FH_ cells, T_PH_ cells, germinal center inhibition, vaccine responses, autoimmunity

## Abstract

Obesity is a rapidly growing public health crisis associated simultaneously with increased metabolic disease and humoral immune suppression to vaccination or infection. Inflammatory CD11c+T-bet+ B cells increase in spleen and adipose tissue during obesity and exacerbate metabolic dysfunction via antibodies. We now find that during obesity Tbet+ B cells also expand in the liver but not omentum or mesenteric fat. Obese mice also develop increased splenic CXCR5+ T_FH_ and hepatic CXCR5-T_PH_ cells which serve as likely partners for antigen-experienced MHC-II+ CD11c+ Tbet+ B cells. We also observed that antibodies in obese mice, previously found to contribute to metabolic disease, largely circulate as inflammatory autoantigen-bound immune complexes. Obese mice lacking T-bet in B cells also develop increased autoantibody titers and expanded splenic germinal center (GC) B and T helper cells. T-bet+ B cell-deficient mice make a similarly enhanced GC, T_FH_, T_PH_ response to haptenated-protein vaccination with a corresponding increase in antibody affinity, although there is no additive effect of obesity. These results are consistent with GC inhibition by expanded CD11c+ B cells demonstrated by others to occur during autoimmunity, suggesting a broadly universal mechanism which may explain reduced humoral immunity and poor clinical outcomes following infection in patients with obesity and other forms of chronic inflammation.

## INTRODUCTION

Obesity is a growing public health crisis associated with many comorbidities including cardiovascular disease, NAFLD, type 2 diabetes, respiratory infections, and cancers such as hepatocellular carcinoma ^1^. Obesity is also a major risk factor for premature death, poor clinical outcomes, and excess healthcare spending across the world. During the recent COVID-19 pandemic, it become evident that obesity also compromises the ability of patients to mount protective antibody responses during infection and in response to vaccination ^2,3^. Following SARS-CoV-2 infection, obese patients present with antibodies that initially reach similar titers as those found in lean patients, but the titers in obese patients wane more quickly and have lower neutralization capacity than antibodies in lean patients^3–7,8^. The majority of antibodies in SARS-CoV-2 infected obese patients may not even be pathogen-specific antibodies, but unwanted autoimmune antibodies ^9^. The mechanism of obesity-induced impairment of protective antibodies is unknown but could result from a skewing of activated B cells away from germinal center responses and towards extracellular reactions ^10^, an active inhibition of GCs ^11^, or alterations in supportive T helper cells. We recently reported increased frequencies of highly-activated CD11c+T-bet+ B cells within the subcutaneous adipose tissue of overweight and obese patients, as well as the perigonadal adipose tissue and spleens of mice on high-fat diet (HFD) ^12^. While B cells expressing CD11c and Tbet were first identified as a member of a broader phenotypically- and functionally-distinct class of B cells associated with the persistent proinflammatory state of aging (earning the name Age Associated B cells: ABCs) ^13,14^, cells with similar or related phenotypes are also recognized to arise during various other forms of prolonged immune activation and may preferentially reside in the extrafollicular space. B cells with a similar immunophenotype (described as “atypical memory” or “double-negative 2” [DN2] B cells) have been found to accumulate in an assortment of autoimmune diseases ^15–26^, infections ^27–34^, and the chronic inflammation of obesity ^12,35^. Deletion of T-bet in B cells significantly reduces weight gain and improves glucose tolerance in HFD mice, indicating a pathogenic role for these cells in the development of obesity-related metabolic disease ^12^.

Studies with autoimmunity models have shown that expansion of CD11c+T-bet+ B cells depends upon innate engagement via TLR7/9 in the context of T cell help via Th1 cytokine or costimulatory signals ^13,15,36^. Cytokine requirements for T-bet+ B cell expansion include IL-21 and IFNγ, which promote B cell expression of CD11c and T-bet, respectively ^25,36–38^. Evidence of somatic hypermutation ^39^ and secretion of class-switched IgG2a/c antibodies by these cells ^31,40^ suggests that CD11c+T-bet+ B cells require help from CD4+ T cells to expand. This is further supported by evidence that B cell expression of MHC class II, CD40, and CD40L were also necessary for CD11c+T-bet+ B cell accumulation during aging ^39^.

However, Song and colleagues ^41^ have observed that during viral infection, splenic CD11c+T-bet+ B cells arise prior to and independently of GC responses and conventional T help, instead depending on extrafollicular interactions with T helper cells. Furthermore, Levack et al. ^38^ have demonstrated that deletion of conventional Bcl-6-dependent T_FH_ cells does not impair the formation of T-bet+ B cells during E. muris infection, an infection characterized by disrupted splenic GC architecture. Instead, during *E. muris* infection, genetic ablation of T-bet-expressing CD4+ T cells, including a Th1-polarized subset of CXCR5+ T_FH_ cells (T_FH_1) and population of CXCR5-T_FH_-like cells, was associated with diminished CD11c+T-bet+ B cell responses ^38^. These data suggest CD11c+Tbet+ ABCs may receive extrafollicular help from CD4+ helper T cells secreting IFNγ and IL-21 and expressing CXCR3 ^38,42^, which promotes migration towards peripheral sites of inflammation ^42,43^, but not expressing CXCR5, which promotes migration into the GC. Among the growing list of T_FH_-like cells that are implicated in driving extrafollicular B cell responses are the PD-1^HI^ CXCR5-Bcl-6-ICOS+ CD4+ T peripheral helper (T_PH_) cells, which share many of the same B-helper effector functions as T_FH_ cells but localize primarily in chronically inflamed peripheral tissues rather than GCs ^44–46^. T_PH_ cells have been found to correlate with atypical memory B cell frequencies in autoimmunity ^25,47–49^, and infection ^25,50^, and both T_FH_1 and T_PH_ cells have been demonstrated to drive T-bet expression in memory cells in vitro ^25,48^. However, whether T_FH_1 cells and/or T_PH_ cells contribute to the HFD-induced expansion of T-bet+ B cells during obesity remains untested.

In this study, we determine that many of the serum IgG autoantibodies in obese mice exist in the form of circulating IgG immune complexes, containing autoantigens such as DNA and RNA. The immune complexes arise more prominently in female mice, consistent with the female sex-bias found during most autoimmune diseases. During obesity, we observe that IC levels correlate with increased levels of CD11c+ T-bet+ B cells within the liver and spleen, but not the mesenteric adipose tissue or omentum. In addition to CD11c+Tbet+ B cells, we note that obese mice develop increased frequencies and numbers of PD-1^HI^ ICOS+ CXCR3+ CD4+ T cells within the liver (T_PH_ cells) and spleen (T_FH_ cells). Both T helper subsets also simultaneously secrete IL-21 and IFNγ, and express CXCR3, fulfilling many of the criteria needed by helper cells for CD11c+Tbet+ B cells. To complement T cell expression of CXCR3, CD11c+ Tbet+ B cells produce CXCL10 ^12^ and express antigen presenting molecules such as MHC II and CD86. Finally, we determined that obese mice lacking T-bet in their B cell compartment have a more pronounced splenic GC response, characterized by increased splenic class-switched B cells, antigen-specific B cells and T_PH_ cells. Interestingly, this is not restricted to obesity-induced autoantigen-specific responses, since obese animals lacking T-bet+ expression in B cells also develop expanded humoral GC B cell responses following haptenated-protein immunization. Thus, obesity-induced expansion of CD11c+T-bet+ B cells skews immune responses away from GC responses or inhibits GC responses directly, resulting in reduced B cell and antibody responses to autoimmune and vaccine antigens. These results may help to explain the reduced effectiveness of B cell responses to pathogens and vaccination in individuals with expanded populations of CD11c+Tbet+ B cells, including those with obesity.

## RESULTS

### T-bet+ CD11c+ B cells expand in spleen, liver, and adipose tissue of obese mice

To explore the effect of obesity on antibody responses, we first optimized our diet-induced obesity (DIO) model using female C57BL/6 (WT) mice. Young female mice can be inconsistently resistant to obesogenic effects of HFD ^51,52^, so initial studies compared the effects of ad libitum HFD-feeding in female mice starting at either 8 or 16 weeks-of-age. Both approaches resulted in significant weight gain in HFD mice compared to age-matched NCD controls (**Fig S1A-D**), with female mice starting HFD diet at both 8 and 16 weeks-of-age exhibiting consistent and significant increases in total body weight as well as WAT, omental, and MAT mass (**Fig S1E-G**). Female mice fed a HFD starting at 16 weeks-of age exhibited significant increases in leukocyte number in spleen, liver, WAT, omentum, and MAT while female mice starting their HFD earlier at 8 weeks only developed significant increases in leukocyte number in WAT and MAT, not the other tissues (**Fig S1H-L**).

We next compared splenic leukocyte population subsets in female mice fed a HFD starting at 8 vs 16 weeks of age (**Fig S2**). Flow cytometry revealed relative (percentages) and absolute (total numbers) obesity-induced increases in the B cell compartment of female mice starting HFD at 16 weeks vs 8 weeks (**Figure S2A,K**). We next used CD21- and CD23-specific antibodies to compare the percentages and numbers of follicular (FO, CD21+ CD23+), marginal zone (MZ, CD21^high^ CD23-), and conventionally-defined age-associated B cells (ABC, CD21-CD23-) in both groups of female mice (**Figure S2B-D, G-I, L-N, Q-S**). In both age groups, HFD mice had a higher number of FO B cells than lean NCD-fed counterparts, but their relative frequencies did not change in the 8 week group and were only modestly increased in the 16 week group, showing that within those mice the FO B cell population expanded proportionally with the total B cell compartment expansion (**Figure S2B,G,L,Q**). On the other hand, in both age groups the percentage of MZ B cells was significantly decreased in HFD mice, while their absolute numbers remained constant, suggesting that this population remains unchanged with obesity resulting in their relative decrease when other populations expand (**Figure S2C,H,M,R**). CD21-CD23-ABCs were increased in number in groups fed a HFD starting at either 8 or 16 weeks, but the increase only reached significance in the mice fed HFD at 16 weeks (**Fig S2D,I,N,S**). Interestingly, the absolute numbers of CD11c+Tbet+ B cells were increased in both 8 and 16 week groups of HFD mice, and the relative percentages were also increased in both sets of mice but only reached significance in the mice starting the HFD at 8 weeks (**Figure S2F,L**). Looking closely reveals that the lack of significant differences between the 16 week NCD and HFD groups most likely reflects an age-related increase in the background of CD11c+Tbet+ B cells in NCD control mice. A broad survey of tissue specific changes in mice fed a HFD starting at either 8 weeks and 16 weeks (both sets of mice pooled) confirmed that, in general, CD11c+T-bet+ B cells increased in number in the spleen, WAT, and liver but not omentum or MAT following HFD feeding (**Fig. 1A-E**). Mice fed a HFD starting at 16 weeks of age will be used for all obese animals in the remainder of the studies. Because the ABC subset (as defined by CD21-CD23-B cells) also includes subsets of cells that are T-bet^neg^ or CD11c^neg^, we chose the T-bet+ CD11c+ B cell population for all remaining analyses.

**Figure 1.**
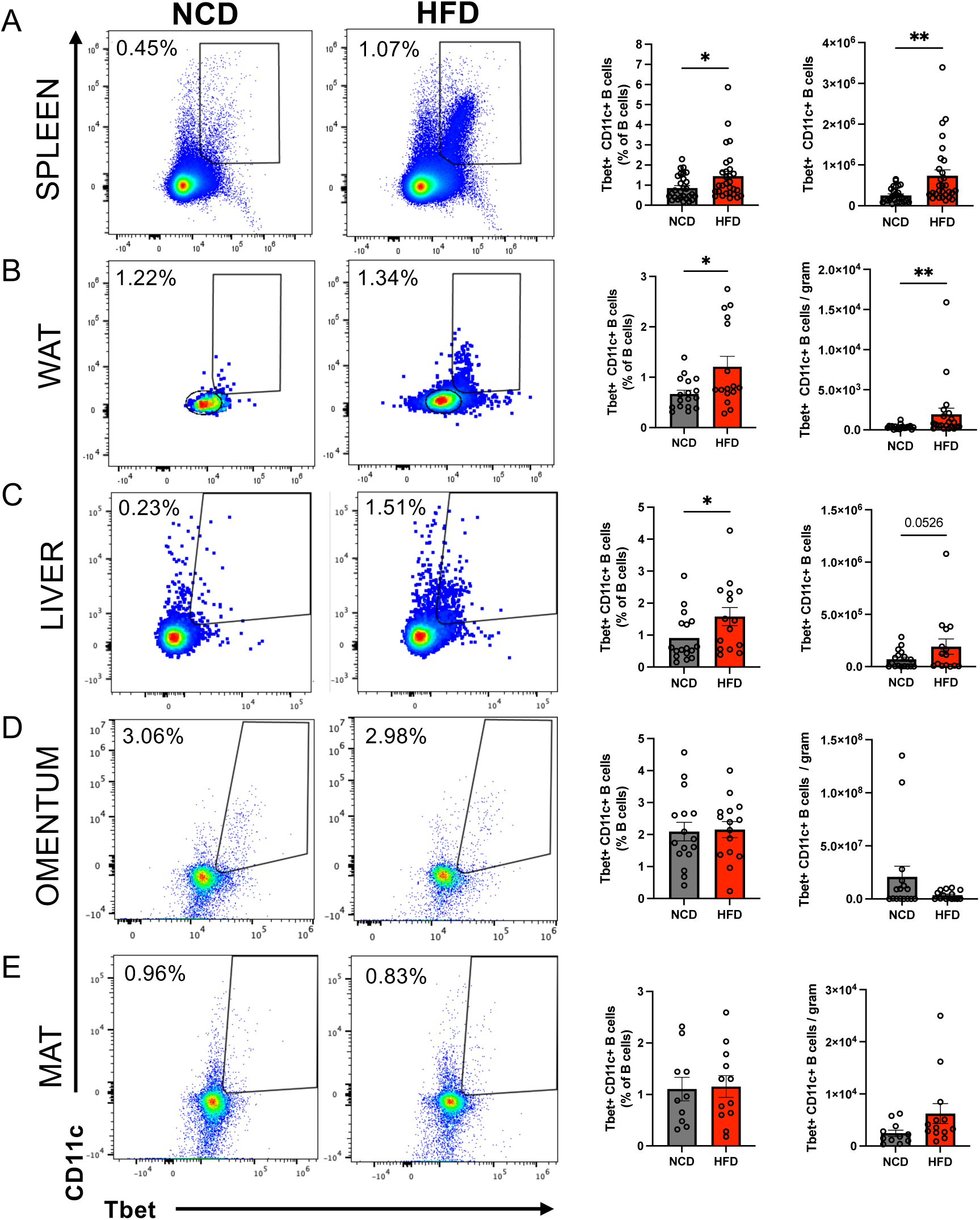
T-bet+ CD11c+ B cells increase in spleen, perigonadal fat (WAT), and liver, but not visceral depots such as omentum or mesenteric fat (MAT) in HFD-fed female mice. (A-D) Pooled analysis of female WT mice starting diet at 8 weeks-of-age and 16 weeks-of-age. Left most panels depict representative flow cytometric plots of B220+ CD19+ CD11c+ T-bet+ B cell frequencies in spleen (A), WAT (B), liver (C), omentum (D), and MAT (E) of either NCD (left) or HFD (right) - fed mice. Right most panels include bar graphs summarizing frequency and number of B220+ CD19+ CD11c+ T-bet+ B cells in individual mice from NCD (black) and HFD (red) - fed groups. Circles represent individual mice, data pooled from 2 independent experiments per age group, n= 4 mice/group. Bar =mean ± SEM; [Student’s t-test (two-tailed) (A-D)]. *p < 0.05, **p < 0.01.

### High levels of circulating immune complexes in obese female mice depend on Tbet+ B cells

Given the broad increases in CD11c+Tbet+ B cells across different peripheral tissues during obesity, and previous evidence for IgG-mediated exacerbation of metabolic disease ^53^, we next examined the humoral B cell contribution to inflammation during obesity. We compared levels of circulating immune complexes (ICs) using ELISA plates coated with C1q molecules, a complement protein which binds to antigen-bound antibodies with a higher affinity than un-bound antibodies. IgG IC concentrations in serum were significantly increased in mice fed a HFD (**Fig 2A, left panel**), and the levels of IgG ICs correlated significantly with the weight of the mice (**Fig 2A, right panel**). This correlation was evident in both males and females but was more profound in female mice (**Fig 2A, right panel**). IgM ICs were not altered in either female or male mice with obesity, nor did the concentrations correlate with animal weight (**data not shown**). Similar to effects observed during autoimmunity, the IgG ICs also deposit in the kidney glomeruli of mice fed a HFD but not in lean NCD-fed mice (**Fig 2B**). The dominant isotypes of ICs in HFD-fed mice were IgG1 and IgG2c ICs (with an upward trend for IgG2b) as compared to NCD-fed lean mice (**Figure 2C**). On the other hand, IgG3 IC levels did not change regardless of the diet (**Figure 2C**).

**Figure 2.**
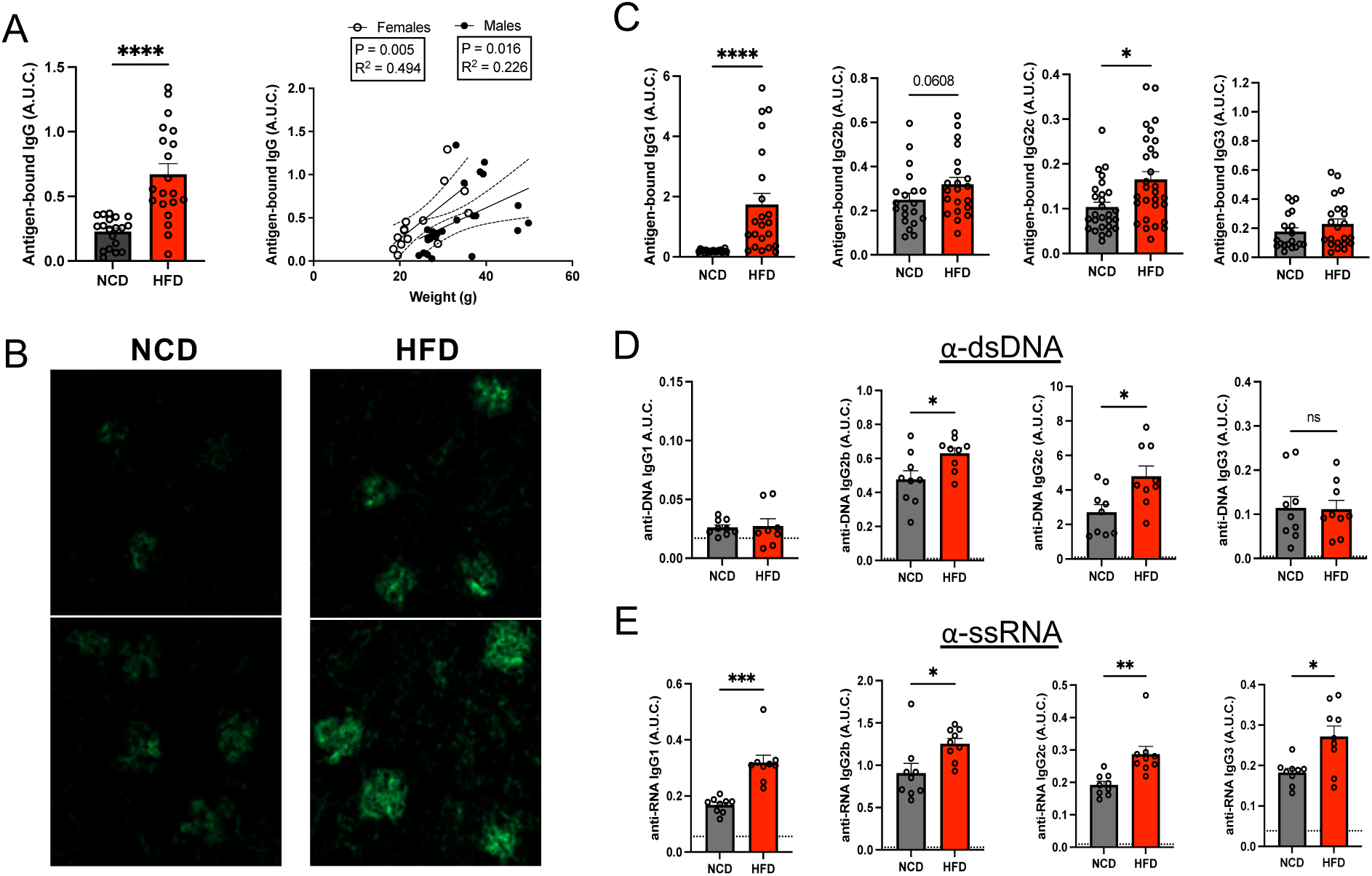
HFD fed-obese mice develop increases in serum Immune Complexes, especially IgG1 and IgG2c, including anti-DNA and RNA autoantibodies when compared to NCD-fed lean controls. C1q ELISA reveals levels of IgG antigen-bound antibodies (immune complexes) in NCD and HFD-fed female mice (**A, left panel**) and allows comparison of correlations between antigen-bound IgG and mouse weight between female and male mice fed a HFD (**A, right panel**). Immunofluorescent labeling of 6μm kidney sections from NCD and HFD WT mice identifies IgG deposits in glomeruli in NCD and HFD-fed female mice (**B**). Levels of serum antigen-bound IgG1, IgG2b, IgG2c, and IgG3 antibody immune complexes, as measured by C1q ELISA, in NCD and HFD WT female mice (**C**). Antigen-specific ELISA reveals anti-dsDNA (**D**) and anti-ssRNA (**E**) levels for IgG1, IgG2b, IgG2c, and IgG3 antibody from NCD (grey) and HFD (red) - fed female mice. Data is representative (**B**) or a pool (**A, C-E**) of 2-4 experiments, 2-10 mice per group, 18-25 weeks old. Bars indicate mean ± SEM. Each symbol indicates an individual mouse. [Non-parametric Unpaired two-tailed t-test (**A**), Two-way ANOVA (**C-E**), and Simple linear regression (**A**)] *p ≤ 0.05, **p ≤ 0.01, ***p ≤ 0.001, ****p ≤ 0.0001

Previous studies have identified antibodies elevated during obesity to be autoreactive ^54^, and the detection of ICs during sterile inflammation suggested self-antigens were available and being engaged, so we measured levels of autoantigen-specific antibodies in obese mice. Autoantibodies in serum of obese mice likely cover a range of specificities, but anti-DNA ELISA revealed significantly increased IgG2b and IgG2c anti-dsDNA antibodies in HFD mice compared to lean mice (**Figure 2D**). Additionally, all four isotypes of anti-ssRNA antibodies were elevated in the HFD mice compared to NCD lean controls (**Figure 2E**). CD11c+Tbet+ B cells typically produce IgG2c antibodies, so to assess the contribution of CD11c+Tbet+ B cells to circulating ICs during obesity, we next compared serum from NCD and HFD - fed mice lacking Tbet in the B cell population (**Fig 3**). To create these mice, we crossed CD19^Cre^ mice with Tbx21^loxP/loxP^ or Tbx21^+/+^ mice to generate knock-out (KO) mice where Tbx21, the gene encoding T-bet, was silenced only in B cells (T-bet+ B KO: B^TbetKO^) and appropriate intact CD19^Cre^ littermate controls (T-bet+ B LC: B^WT^). As expected, B^TbetKO^ mice developed lower amounts of IgG2c ICs than B^WT^ controls regardless of diet (**Fig 3C**). IgG1 ICs elevated in HFD-fed B^WT^ mice compared to lean NCD controls were also dependent upon Tbet expression in B cells **(Fig 3A**). In absence of IgG1 and IgG2c ICs, levels of IgG3 ICs were elevated in B^TbetKO^ mice regardless of diet, although only reached significance in lean mice (**Fig 3D**). IgG isolated from HFD-fed obese mice has been previously found to exacerbate metabolic disease^12^, so we next used in vitro culture of adipose stromal vascular fraction leukocytes to determine whether the purified ICs from lean or obese mice were similarly capable of inducing inflammatory cytokines TNFalpha and IL-6 (**Fig 3E**). We determined that equal concentrations of ICs from lean and obese mice induced similar levels of TNFalpha but the ICs from HFD mice stimulated more IL-6 than NCD-fed mice suggesting there are unique differences (**Fig 3F**).

**Figure 3.**
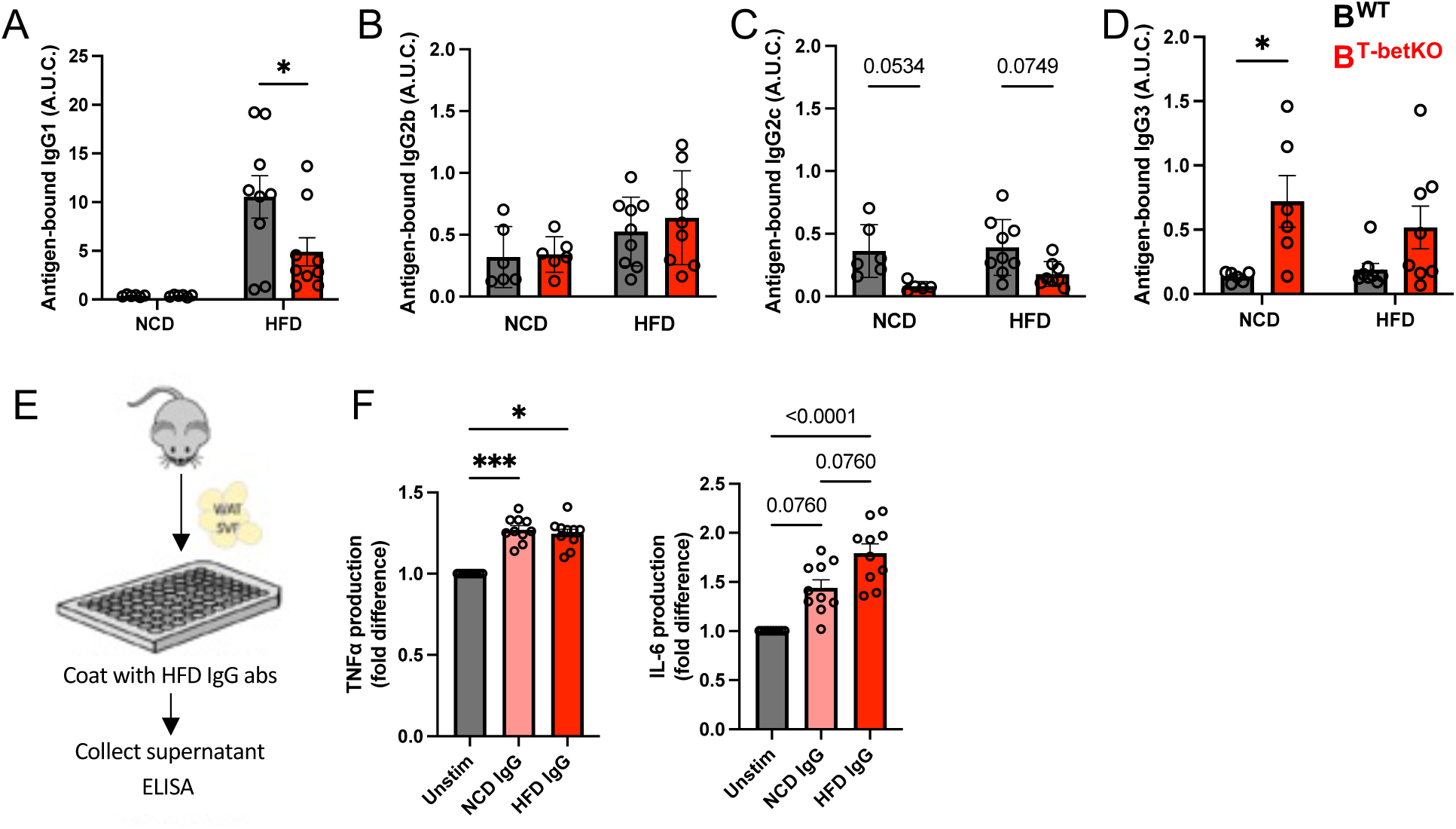
IgG1 and IgG2c serum Immune Complexes (ICs) depend on Tbet+ in B cells. IgG ICs can drive in vitro inflammatory cytokine production from stromal vascular fraction leukocytes. Levels of serum antigen-bound IgG1 (**A**), IgG2b (**B**), IgG2c (**C**), and IgG3 (D) antibody immune complexes, as measured by C1q ELISA, were detected in NCD and HFD-fed B^WT^ littermate controls (grey) and B^TbetKO^ (red) female mice. In vitro stimulation of stromal vascular fraction (SVF) leukocytes from WAT with 31ug column purified IgG from NCD and HFD-fed mice (**E**). 24hr supernatants were evaluated by cytokine ELISA for TNFa and IL-6 (F). Data is a pool of 2 experiments, 4-5 female mice per group, 18-25 weeks old. Bars indicate mean ± SEM. Each symbol indicates an individual mouse. [One-way ANOVA] *p ≤ 0.05, **p ≤ 0.01, ***p ≤ 0.001

### Obese mice have expanded splenic GC B and ASC

Class-switched B cells responding to conventional T-dependent antigens typically develop in the context of a germinal center (GC) where the B cells localize for a carefully orchestrated interaction which allows them to receive T cell help, undergo affinity maturation, and class switch. To consider a source for the autoantigen specific, class-switched B cells producing IgG for ICs in mice fed a HFD, we assessed frequency and numbers of splenic germinal center B cells (GC, CD95+ GL7+) and antibody-secreting cells (ASCs, CD138^high^ IgD-) in the context of total B cells). HFD-fed mice show increases in total splenocytes as well as increases in absolute number and frequency of B cells (**Fig 4A**). Both GC B cells and ASCs had specifically expanded numbers, but their percentages were not significantly increased, which is consistent with the general enlargement of the B cell compartment (**Figure 4B**). Increases in numbers of antigen-experienced B cell populations such as the GC and ASCs tested here, as well as the CD11c+Tbet+B cells tested in **Fig 1**, are consistent with the chronic inflammation and ongoing autoimmunity detectable in obese mice. Furthermore, the CD11c+T-bet+ B cells in the spleen are well suited to engage with and present antigen to T cells, as they express higher levels of CD86 and MHC-II than follicular or marginal zone B cells, regardless of diet (**Fig 4C,D**). Obese patients and mice develop high levels of free fatty acids (FFA) in serum, which can be cytotoxic, so we next tested the consequences of high concentrations of extracellular FFA, on CD11c+T-bet+ B cells. To test this, we activated total splenocytes from NCD mice *in vitro* using a cocktail of anti-CD40, R848, IL-21, and IFNγ, in combination with increasing concentrations of oleate (OA, unsaturated fatty acid), palmitate (PA, saturated fatty acid), or control BSA. High concentrations of palmitate induced significant cell death in the B cells after three days of culture (**Figure S3A**). This result confirmed previous literature on the lipotoxic effects of saturated fatty acids ^55,56^. The percentages of T-bet+ B cells in our cultures dramatically increased in the palmitate-treated group and the absolute numbers of T-bet+ B cells remained constant despite the significant amount of cell death among total B cells (**Figure S3B**). Together, these results suggest that T-bet+ B cells are able to withstand lipid-induced stress and cell death to a much higher degree than other splenic B cells.

**Figure 4.**
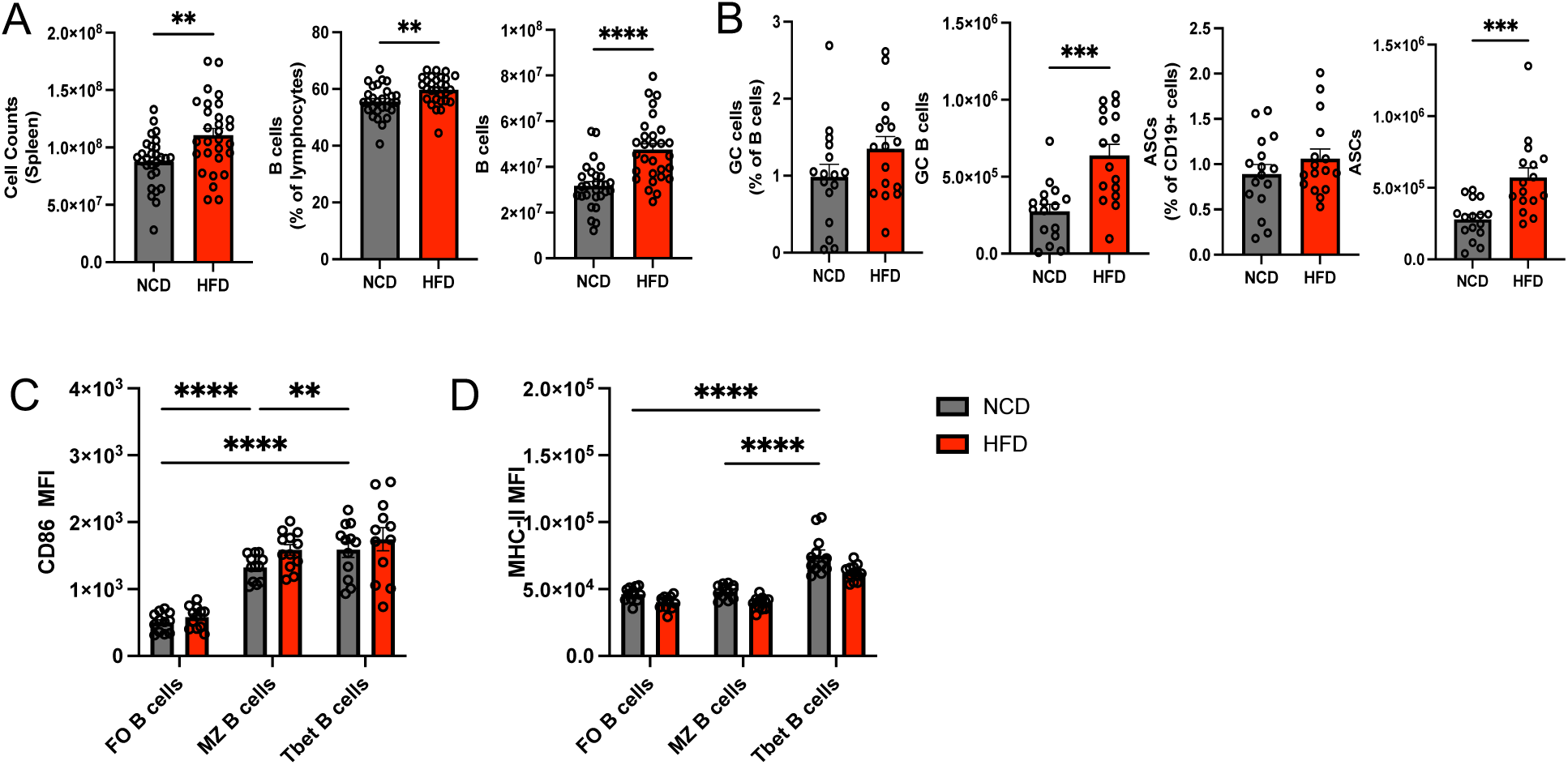
Obese mice fed a HFD develop increased number and frequency of splenocytes and B cells, with specific increases in number of GC B cells and ASCs. CD11c+Tbet+ B cells express higher levels of CD86 and MHCII than other B cell subsets, regardless of diet. Cell counts as well as B cell frequencies and absolute numbers of splenocytes from NCD and HFD WT mice (**A**). Summary of frequencies and absolute numbers of CD19+ GL7+ CD95+ germinal center B cells and CD19+ IgD+ CD138+ antibody secreting cells (ASC) (**B**) in the spleen of NCD and HFD WT mice. Flow cytometry detection of CD86 (**C**) and MHC-II (D) expression, in splenic B cell subsets from NCD and HFD WT female mice. Data is a pool of 4-7 experiments, 4-5 female mice per group, 21-30 weeks old. Bars indicate mean ± SEM. Each symbol indicates an individual mouse. One data point was excluded based on ROUT test (Q = 0.1%). [Non-parametric Unpaired two-tailed t-test (**A,B**); 2way ANOVA(**C,D**)] *p ≤ 0.05, **p ≤ 0.01, ***p ≤ 0.001, ****p ≤0.0001

### Obese mice have increased number of class-switched CD11c+Tbet+ B cells in adipose tissue

We next asked if leukocytes in adipose tissue of obese female mice started on their HFD at 16 weeks displayed changes similar to those observed in splenocytes from the same animals. Indeed, leukocytes in the Stromal Vascular Fraction (SVF) from HFD fed animals developed a similar increase in absolute number of total B cells and CD11c+ T-bet+ B cells as was observed in the spleen, when compared to NCD fed controls (**Figure S4A-D**). Frequency of total B cells, but not frequency of CD11c+Tbet+ B cells, was increased suggesting the increased numbers of CD11c+Tbet+ B cells reflected a general B cell increase (**Figure SA,C**). Staining for surface immunoglobulin expression on adipose tissue B cells (**Figure S4E,F**) revealed that the T-bet+ B cells represent a highly class-switched (IgM-IgD-) subset (**Figure S4G**). In HFD-fed mice, the class-switched T-bet+ B cells amount to nearly half of the total class-switched adipose B cells. Adipose tissue CD11c+T-bet+ B cells also exhibited similar high expression levels of proteins consistent with activation and proliferation, including CD86 and Ki-67 (**Figure S4H-J**) as was observed for CD11c+Tbet+ B cells in the spleen (**Fig 4C).** However, adipose CD11c+Tbet+ B cells and Tbet-B cells do not express different levels of MHCII in the adipose tissue (**Fig S4I**) the way they do in the spleen (**Fig 4D**), suggesting interactions with helper T cells may be similar for all B cells in the adipose tissue. CD11c+Tbet+ B cells in adipose tissue also express chemokine receptors CXCR4 and CXCR5 but not CXCR3 (**Fig S4K-M**), regardless of diet. CD11c+Tbet+ B cells in adipose also express elevated levels of scavenger receptor CD36 and show high levels of intracellular neutral lipids, regardless of diet (**Fig SN,O**). This suggests that HFD alone does not dictate adipose localization and phenotype of CD11c+Tbet+ B cells. Instead, any CD11c+Tbet+ B cells that make it into adipose tissue appear to be activated and prepared to interact with helper cells, regardless of diet; there are just more of them in mice fed a HFD.

### Obesity expands IFNγ+IL-21+CXCR3+ T_FH_ and T_PH_ cells in the spleen and liver

T-bet+ B cells are class switched and express antigen presenting and costimulatory molecules which positions them for productive interactions with CD4+ T_H_ cells ^25,38,39,41^. Total CD4+ T cell frequency and number are the same in spleens from NCD and HFD-fed mice but are significantly expanded within the liver (data not shown). We examined the T cells for co-expression of PD-1 and C-X-C chemokine receptor 5 (CXCR5), markers used to identify specialized B-helper CD4+ T cell subsets ^57–61^. We found that HFD preferentially expanded the frequency and number of CXCR5 ^pos^ PD-1^HI^ CD4+ T cells in the spleen [**Fig. 5A (top, blue gates);** quantified in **5B**] and CXCR5^neg^ PD-1^HI^ CD4+ T cells in the liver [**Fig. 5A (bottom, yellow gates);** quantified in **5C].** Subsequent analysis of these populations confirmed the identity of the CXCR5 ^pos^ PD-1^HI^ CD4+ T cells in both spleen and liver as bona fide T_FH_ cells, which expressed elevated levels of the lineage-defining T_FH_ transcription factor B cell lymphoma 6 (Bcl-6) (**Fig. 5D**) ^61–64^, and the inducible T cell co-stimulator (ICOS) (**Fig. 5E**). In contrast, the PD-1^HI^ CXCR5^neg^ CD4+ T cell population in both tissues lacked Bcl-6 expression (**Fig. 5D**) but expressed ICOS (**Fig. 5E**) at levels comparable to the CXCR5 ^pos^ T helper cells, consistent with identity as T peripheral helper (T_PH_) cells.’, Th1-polarized T_FH_ (T_FH1_) cells co-express Bcl-6 and T-bet ^42,65^, and are important for directing B cell antibody class-switching to IgG2a/c ^42,66^, an isotype favored by murine CD11c+T-bet+ B cells ^14,31^, so we next tested T cells for T-bet expression. Both CXCR5 ^pos^ and CXCR5^neg^ T helper cells express the transcription factor T-bet (**Fig. 5F**). Thus, a HFD primarily expands a CXCR5+ T_FH_ population of PD-1^HI^ CD4+ T cells in the spleen and a CXCR5-T_PH_ population of PD-1^HI^ CD4+ T cells within the liver, indicating that distinct PD-1^HI^ CD4+ T cell populations may support T-bet+ B cell expansion within different anatomic niches.

**Figure 5.**
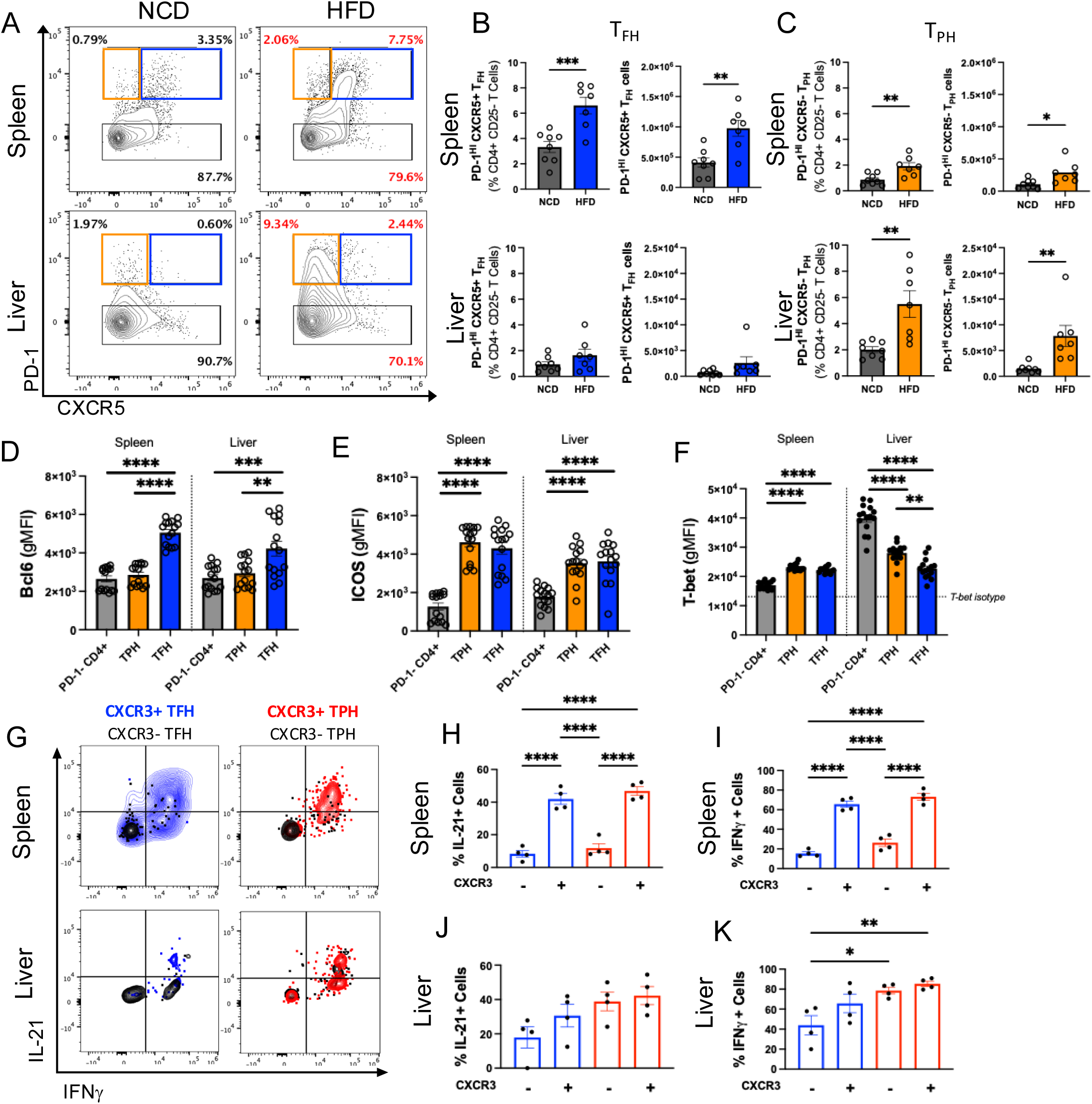
Obese mice have expanded populations of CXCR3+, IFNg+, IL-21+ T_FH_ cells in spleen and T_PH_ cells in liver. Representative flow cytometry contour plots of PD-1 x CXCR5 expression within CD25-CD4+ T cells isolated from NCD (left) and HFD (right) mouse spleen (top) and liver (bottom) (**A**). Blue gates represent PD-1^HI^ CXCR5+ cells, yellow gates represent PD-1^HI^ CXCR5-cells, and black gates represent PD-1-cells (**A**). Frequency and absolute number of PD-1^HI^ CXCR5+ T_FH_ cells (**B**, blue), or PD-1^HI^ CXCR5-T_PH_ cells (**C**, yellow) within the CD25-CD4+ T cell compartments of spleen (top) and liver (bottom) of NCD (grey) and HFD (blue/yellow) mice. Quantification of Bcl-6 (**D**), ICOS (**E**), and T-bet (**F**) protein expression (gMFI) in splenic (left) or hepatic (right) CD25-CD4+ T cell subsets. Representative flow cytometry contour plots depicting relative expression of IL-21 and IFNg by PD-1^HI^ CXCR5+ T_FH_ cells or PD-1^HI^ CXCR5-T_PH_ cells from spleen (top) or liver (bottom) that are also CXCR3+ (blue/red) or CXCR3-(black) (**G**). Summary of frequency of IL-21 (**H,J;** left) or IFNg (**I,K**; right) producing CD4+ PD-1^HI^ CXCR5+ T_FH_ (blue) or PD-1^HI^ CXCR5-T_PH_ (red) cells categorized by CXCR3+ expression as noted in spleen (**H, I**) or liver (**J,K**) from HFD-fed mice. Data pooled from 2 independent experiments with 4 female mice/group, Bar ± SEM; [Student’s t-test (two-tailed) (B,C); one-way ANOVA (D-F; H-K)]; *p < 0.05, **p < 0.01, ***p < 0.001, ****p < 0.0001

The cytokines IFNγ and IL-21 are chiefly responsible for CD11c+T-bet+ B cell generation ^25,36–38^, and T_PH_ and T_FH1_ cells can both serve as sources for these cytokines ^25,38,42,44,48,65,67,68^. Given the strong correlation between the expanded populations of PD-1^HI^ CD4+ T helper cells and T-bet+ B cells we observed within the HFD liver and spleen, we next assessed the production of IFNγ and IL-21 from T cells isolated from both tissues using flow cytometry (**Fig. 5G**). Very few T_FH_ or T_PH_ cells are detectable in NCD animals, so we restricted all subsequent analysis to these populations in HFD-fed animals.

Intracellular cytokine staining revealed cytokine-secreting CXCR5+ T_FH_ and CXCR5-T_PH_ cells were present as IFNγ/IL-21 double-positive and IFNγ single-positive populations (**Fig. 5G**). Comparison of IFNγ and IL-21 production across T cell subsets in the HFD liver and spleen showed that within both tissues, PD-1^HI^ CD4+ T cells were the predominant producers of IL-21 and tended to also be making IFNγ (**Fig. 5G**). Intracellular cytokine staining of splenic cells stimulated ex vivo indicated that CXCR3+ T_FH_ and T_PH_ cells had higher frequencies of IL-21 as well as IFNγ expression than their CXCR3-counterparts (**Fig. 5H,I**). On the other hand, in the liver, the degree of IFNγ and IL-21 production was equally distributed among CXCR3+ and CXCR3-T_PH_ cells (**Fig. 5J,K)**). This data demonstrates that HFD promotes the expansion of CXCR3+ IFNγ/IL-21 double-positive T_FH_1 and T_PH_1 cells in the spleen, and predominantly T_PH_1 cells in the liver.

### Obesity does not induce iNKT_FH_ cell differentiation

Our previous studies identified an essential role for iNKT cells in facilitating CD11c+ Tbet+ B cell expansion in murine adipose tissue during obesity. Given the robust increases in CD4+ T_FH_ and T_PH_ cells observed during HFD (**Fig. 5**), we next examined the iNKT cell compartment within the liver and spleen for similar diet-related populations **(Fig. S5**). Within the spleen, we did not detect any meaningful changes in total of CD4+ iNKT cell number or frequency detectable after 16 weeks of HFD (**Fig. S5A-D**). Splenic iNKT cells were CXCR5+ but did not express any traditional T_FH_ markers such as PD-1, Bcl-6, or ICOS (**Fig. S5E-H**) although they remained positive for Tbet (**Fig. S5I**). Similarly, HFD decreased total and CD4+ hepatic iNKT cell frequencies, but not numbers (**Fig. S5J-M**). In contrast to splenic CD4+ iNKT cells, hepatic CD4+ iNKT cells expressed little CXCR5 (**Fig. S5N, O**) and also lacked PD-1, Bcl-6, and ICOS (**Fig. S5O-Q**) but remained highly positive for T-bet (**Fig. S5R**). Thus, iNKT cells may facilitate CD11c+ Tbet+ B cell expansion via early IFNγ, but HFD does not promote increases in iNKT_FH/PH_ cells detectable at this late stage in the liver or spleen.

### CXCR3-blockade promotes splenic T_PH_ retention and increases GC activity

Both diet-expanded populations of T_PH_ and T_FH_ cells expressed high levels of CXCR3, and T-bet+ CD11c+ B cells are important source of CXCR3-ligand, CXCL10(33), so we next examined the consequences of CXCR3 blockade on HFD-expanded PD-1^HI^ CD4+ T cells. Anti-CXCR3 blocking antibodies, which also block CXCL10 (102) and anti-CXCL9, were administered by I.P. injection every other day for two weeks in female mice following 12 weeks of HFD-feeding (**Fig. S6A**). CD4+ T cell compartments were then examined in both spleen and liver by flow cytometry. CXCR3-blockade did not affect total splenic CD4+ T cell numbers, but it did significantly reduce total CD4+ T cells in the HFD liver (**Fig. S6B**). Further analysis showed that blocking CXCR3 resulted in an accumulation of T-bet+ CD4+ T cells within the HFD spleen, and diminished numbers of T-bet+ CD4+ T cells in the HFD liver (**Fig. S6C**), suggesting impaired trafficking of these cells to the periphery. In agreement with this interpretation, the spleen exhibited increased amounts of PD-1+ ICOS+ CD4+ T cells (**Fig. S6D**), though this population was unchanged in the liver (**Fig. S6D**). CXCR5 staining was then used to determine the relative changes in T_FH_ and T_PH_ populations in both tissues. As previously seen in HFD mice following genetic ablation of T-bet+ B cells, the rise in PD-1+ ICOS+ CD4+ T cells within the spleen resulted from increased total splenic CXCR5^neg^ T_PH_ cells, rather than total CXCR5 ^pos^ T_FH_ cells (Fig. S6E, F). No changes in either population of PD-1^HI^ CD4+ T cells were observed in the HFD liver (**Fig. S6E, F**). examination of T-bet+ T_FH_ cells in HFD spleens revealed that although total T_FH_ cells were not significantly elevated following CXCR3-blockade, the Th1-polarized T_FH1_ subset was specifically increased (**Fig. S6G**). Congruent with these findings, CXCR3-blockade also led to robust changes in splenic B cells in HFD mice. Specifically, CXCR3-blockade induced greater numbers of splenic B220+ CD19+ GL7+ CD95+ germinal center (GC) B cells (**Fig. S6H**) and GC-derived T-bet+ CD11c+ B cells (**Fig. 6H**). CXCR3-blockade also increased the degree of class-switching in B cells and the number of class-switched T-bet+ CD11c+ B cells (**Fig. 6I**). Therefore, CXCR3-blockade may affect T_FH1_ spatial positioning towards B cell follicles, allowing for a greater degree of positive influence over GC B cells, biasing them towards a T-bet+ B cell immunophenotype.

**Figure 6.**
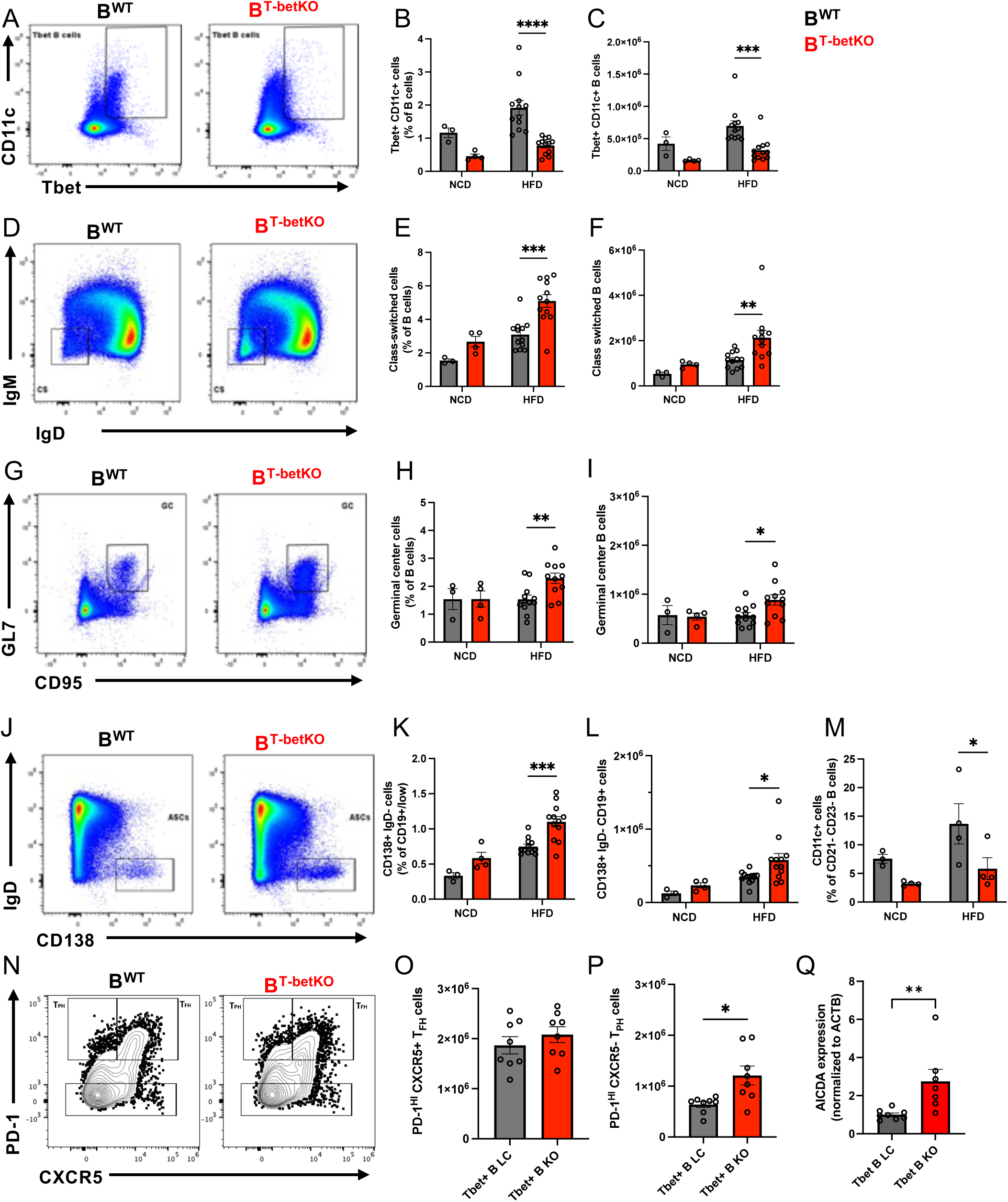
B^TbetKO^ HFD-fed obese mice develop enhanced GC responses with significantly increased class-switched B cells, GC B cells, AICDA expression, antibody-secreting cells, and T_PH_ cells, as well as fewer CD11c+ extrafollicular B cells than intact or NCD-fed lean controls. Representative flow cytometry of specific cell populations in total splenic B cells from B^TbetKO^ and B^WT^ mice paired with summary of frequency and numbers in NCD and HFD-fed B^TbetKO^ (red) and B^WT^ mice (black). Cell populations include splenic CD11c+Tbet+ B cells (**A-C**), IgM- and IgD-class switched B cells (**D-F**), GL7+ CD95+ GC B cells (**G-I**), IgD-CD138+ ASCs (**J-L**). Frequency of CD21-CD23-CD11c+ extrafollicular B cells in spleen of B^TbetKO^ (red) and B^WT^ (black) mice fed a NCD or HFD (**M**). Representative flow cytometry of PD-1 and CXCR5 on CD4+ T cells from B^TbetKO^ and B^WT^ mice (**N**) paired with summary of frequency and numbers in NCD and HFD-fed B^TbetKO^ (red) and B^WT^ mice (black) (**O,P**). Relative AICDA mRNA expression determined by qPCR of isolated splenocytes from HFD B^TbetKO^ and B^WT^ mice (**Q**). Data is a pool of 2 experiments, 4 female mice per group, 27-29 weeks old (**A-P**). Results are from one experiment with 8 female mice per group, 27-29 weeks old (**Q**). One point excluded as an outlier based on ROUT test. Bars indicate mean ± SEM. Each symbol indicates an individual mouse. [Non-parametric Unpaired two-tailed t-test] *p ≤0.05, **p ≤ 0.01, ***p ≤ 0.001, ****p ≤ 0.0001

Despite these changes to the splenic T_FH1_ population, CXCR3-blockade promoted an even greater rise of splenic T_PH1_ cells (**Fig. S6J**). While the liver exhibited a loss in total T-bet+ CD4+ T cells (**Fig. S6C**), no changes were observed in liver T-bet+ PD-1^HI^ populations (**Fig. 6SG, J**). The robust impact on total splenic T_PH_ cells following both genetic ablation of T-bet+ B cells and CXCR3-blockade in HFD mice, suggests that CXCR3-mediates T-bet+ B cell interaction with T_PH_ cells to promote their egress from secondary lymphoid organs.

### Obese mice lacking T-bet expression in B cells exhibit increased autoimmune GC response

Humans with obesity often exhibit reduced humoral responses to infection or vaccination, and Tbet+ ABCs have been suggested to inhibit GC responses during autoimmunity in mice ^69^. To ask if obesity-expanded CD11c+Tbet+ B cells might impose similar restrictions on GC-mediated humoral responses in obese mice, we examined B cell subset alterations induced by HFD feeding in the absence of Tbet-expressing B cells (B^TbetKO^) or intact littermate control mice (B^WT^). We first confirmed that HFD-induced expansion of CD11c+Tbet+ B cells was missing in mice lacking Tbet expression in B cells (**Fig 6A-C**). However, numbers and frequencies of class-switched B cells (**Fig. 6D-F**), germinal center B cells (**Fig. 6G-I**), and CD138+ antibody-secreting cells (**Fig. 6J-L**) were all significantly expanded in the HFD B^TbetKO^ mice compared to HFD B^WT^ controls. In addition to the B cell changes, lean B^TbetKO^ mice exhibited increases in total IgG1 compared with all other mice, while B^TbetKO^ mice fed a HFD developed increases in total IgG2b and IgG3 (**Fig S7A-D**). As expected, B^TbetKO^ mice lacked IgG2c under both lean and obese conditions (**Fig S7C**). Loss of Tbet in B cells in obese mice did not dramatically increase autoantibodies, but we did observe a modest increase in anti-ssRNA of only the IgG2b isotype when B^TbetKO^ mice were fed a HFD (**Fig S7E,F,G**). This is consistent with the possibility that many autoantibodies in this model may not be GC derived, and as such do not increase when GC B cells expand. Since GC B cells increased when Tbet+ was missing from B cells (in B^TBetKO^), we next quantified CD11c+ B cells, considered extrafollicular (EF) B cells, and found a commensurate decrease in EF B cells in B^TbetKO^ mice on a HFD (**Fig. 6M**). In parallel, CXCR5+ T_FH_ cells remained high and CXCR5-T_PH_ cells increased significantly in B^TBetKO^ mice on a HFD (**Fig. 6N-P**), suggesting their regulation may be subsequent to the B cell modifications. Consistent with their increased class switching and enhanced GC response, HFD B^TbetKO^ mice also expressed higher levels of class-switch-driving transcription factor AICDA mRNA than HFD B^WT^ littermate controls, as measured by qPCR (**Fig. 6Q**). These observations show that loss of T-bet expression in B cells during obesity enhances class-switching, germinal center reactions, and differentiation of antibody-secreting cells, supporting the notion that Tbet+ B cells inhibit GC responses in obesity, similar to their effect in lupus ^69^.

### Obese mice lacking T-bet+ B cells exhibit increased GC responses to vaccination

After determining that elimination of T-bet+ B cells enhances autoantibody responses and increases GC B cell populations during obesity, we next asked whether this effect extended to non-self-antigens. The notion that obesity impairs the generation of protective immunity during infection or immunization has been established in several studies ^3–7^. To evaluate the role of T-bet+ B cells in foreign-antigen responses during obesity, we immunized HFD- or NCD-fed B^WT^ and B^TbetKO^ mice IP with NP-KLH and alum to induce a T-dependent response. Despite having a slightly smaller splenic B cell compartment when fed a HFD (**Fig. 7A**), immunized HFD-fed B^TbetKO^ mice developed significantly higher numbers and frequencies of class-switched B cells than immunized HFD-fed B^WT^ controls (**Fig. 7B**)., the immunized B^TbetKO^ mice significantly expanded the number of <u>antigen-specific</u> NP+ B cells (**Fig. 7C**) and number of NP+ class-switched B cells (**Fig. 7D**) as compared to the immunized B^WT^ mice, irrespective of diet. The observation that this antigen-specific effect is diet independent suggests that Tbet+ B cells may be capable of playing the same role in both HFD and NCD mice, but the effect is more evident when inflammation via HFD or immunization with alum activates or expands the B cells.

**Figure 7.**
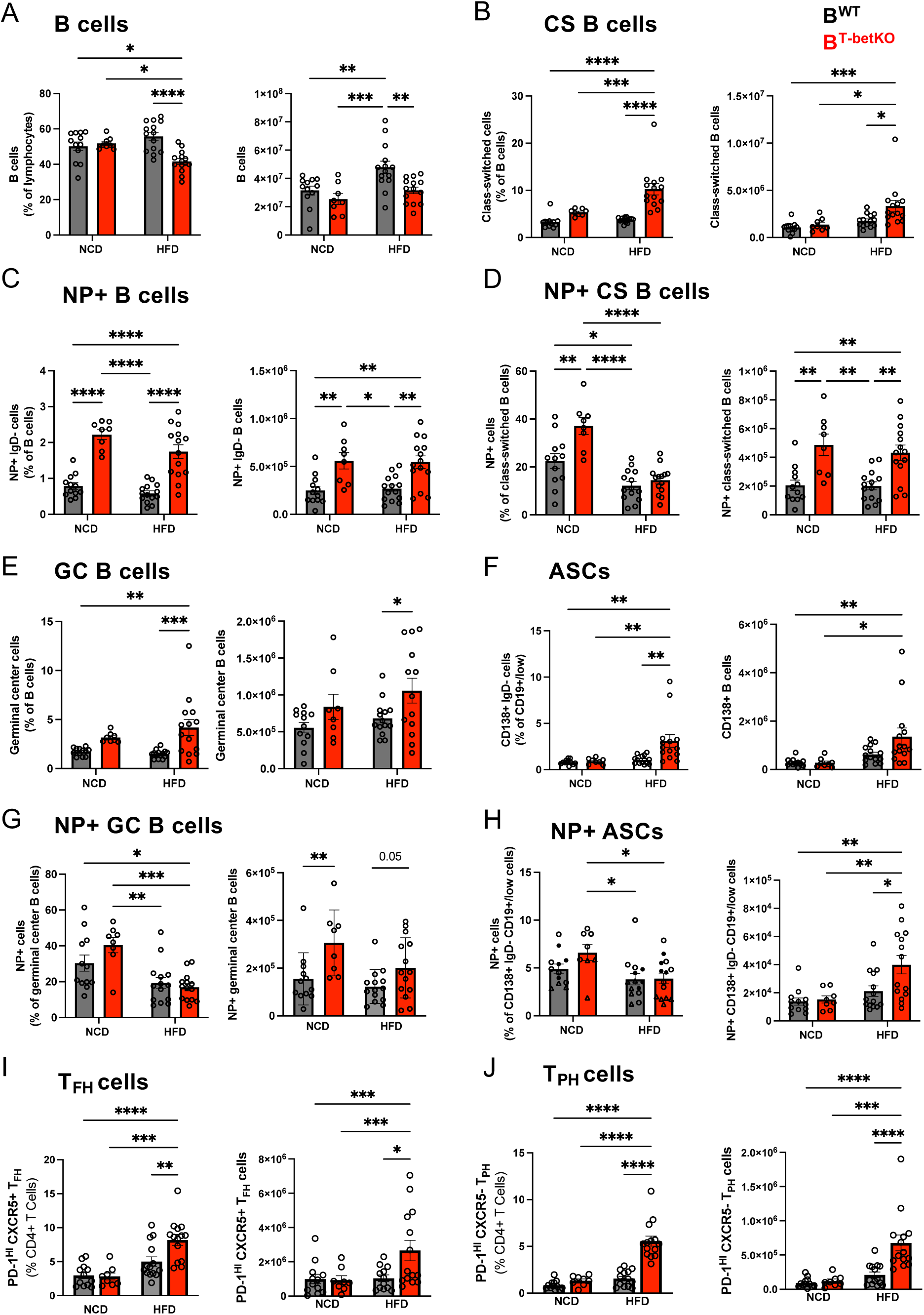
NP-KLH immunized B^TbetKO^ mice mount enhanced GC B cell responses with increased class-switched B cells, GC B cells, ASCs, antigen-specific GCs and ASCs, as well as T_FH_/T_PH_ cells as compared to B^WT^ controls. Summary of flow cytometry measurements of percentages (left) and absolute numbers (right) of spleen lymphocyte populations from NCD and HFD B^TbetKO^ (red) and B^WT^ (black) mice immunized with NP-KLH and alum and assessed 14 days later. Populations include total B cells (**A**), class switched IgM-IgD-B cells (**B**), NP+ B cells (**C**), NP+ class-switched B cells (**D**), GC B cells (**E**), CD138+IgD-antibody secreting cells (ASC) (**F**), splenic NP+ GC B cells (**G**), NP+ ASC (**H**), CD4+PD-1+CXCR5+ T_FH_ cells (**I**), and CD4+PD-1+CXCR5-T_PH_ cells (**J**). Data is pool of two experiments, 4 female mice per group, 30 weeks old. Bars indicate mean ± SEM. Each symbol indicates an individual mouse. [Non-parametric Unpaired two-tailed t-test] *p ≤ 0.05, **p ≤ 0.01, ***p ≤ 0.001, ****p ≤ 0.0001

Immunized HFD-fed, but not NCD-fed, B^TbetKO^ groups also developed an increase in frequency and number of total antigen-non-specific germinal center (GC) B cell populations (**Fig. 7E**) and an increase in frequency plus a trending increase in number of <u>total</u> antigen non-specific antibody secreting cells (ASCs) (**Fig. 7F**) as compared to the B^WT^ groups. This increase in immunization antigen non-specific cells is selective to the HFD groups, suggesting the GCs and ASCs affected may be mediated mostly by obesity-derived antigens rather than the immunization. On the other hand, numbers (but not frequency) of immunization antigen-specific NP+GC B cells (**Fig. 7G**) and NP+ASCs (**Fig. 7H**) were expanded in the B^TbetKO^ as compared to the B^WT^ mice, somewhat irrespective of diet. This mirrors the diet-independent changes observed for immunization antigen-specific NP+ B cells and NP+ CS B cells in (**Fig. 7C**) and (**Fig. 7D**). In sum, this data is consistent with immunization antigen (NP)-specific and obesity antigen-driven GCs and ASCs both increasing in B^TbetKO^ mice, observed as parallel increases in total and immunization antigen-specific B cell number. However, the data also suggests that only the obesity-driven GCs comprise a large enough population to produce changes that reach significance in total frequency in the B^TbetKO^ mice.

Notably, PD-1+CXCR5+ T_FH_ and PD-1+CXCR5-T_PH_ cells were also increased in frequency and number in the immunized HFD-fed B^TbetKO^ mice but not the NCD-fed controls (**Fig. 7I,J**). Thus, following immunization with T-D model antigen and adjuvant, the B^TbetKO^ mice were generally more permissive of an enhanced antigen-specific GC response when compared to their B^WT^ counterparts, irrespective of diet. These data are consistent with evidence from autoimmunity models suggesting Tbet+ B cells inhibit antigen-specific GCs. This relief of inhibition by the loss of Tbet in B cells in B^TbetKO^ mice was only evident for non-NP specific, or total, B cell populations when mice were fed a HFD suggesting these changes may only reach levels of detectability with a combined effect of the immunogen and autoantigens., There were no short-term detectable effects of HFD diet on titer following NP-KLH/alum immunization of B6 WT mice, since NP-specific IgG, IgG1, and IgG2c antibody titers remained the same after primary and secondary vaccination in lean and obese mice (**Fig. 8A**) and there were no specific effect of diet on the increase in NP-specific antibody affinity observed between primary and secondary vaccination (**Fig. 8B**). Since flow cytometry finds increased GC B cells in B^TbetKO^ mice compared to B^WT^ mice (**Fig. 7E**), we next looked for increases in GC size and antibody affinity resulting from removal of Tbet in B cells. These predicted results were confirmed in our lean, NCD-fed mice where affinity of NP-specific IgG from B^TbetKO^ mice was increased compared to intact B^WT^ mice (**Fig. 8C**). However, similar affinity changes were not observed in HFD-fed mice suggesting affinity may already be maximized because of the combined effect of the diet and vaccination with this model hapten/protein (**Fig. 8C**). We next used immunofluorescence to visualize GCs and noted that while general GC structures were similar between the groups, HFD induced increases in GC area in B^WT^ mice compared to NCD-fed controls, as expected. Furthermore, GCs were even larger in B^TbetKO^ mice compared to B^WT^ mice when both are on a HFD (**Fig 8D,E**), which is consistent with earlier flow cytometry data (**Fig. 8E**). These results are also consistent with observations in human patients with obesity who maintain early antibody titers similar to their lean counterparts but still develop reduced neutralization capability or poor clinical outcomes following vaccination or infection.

**Figure 8.**
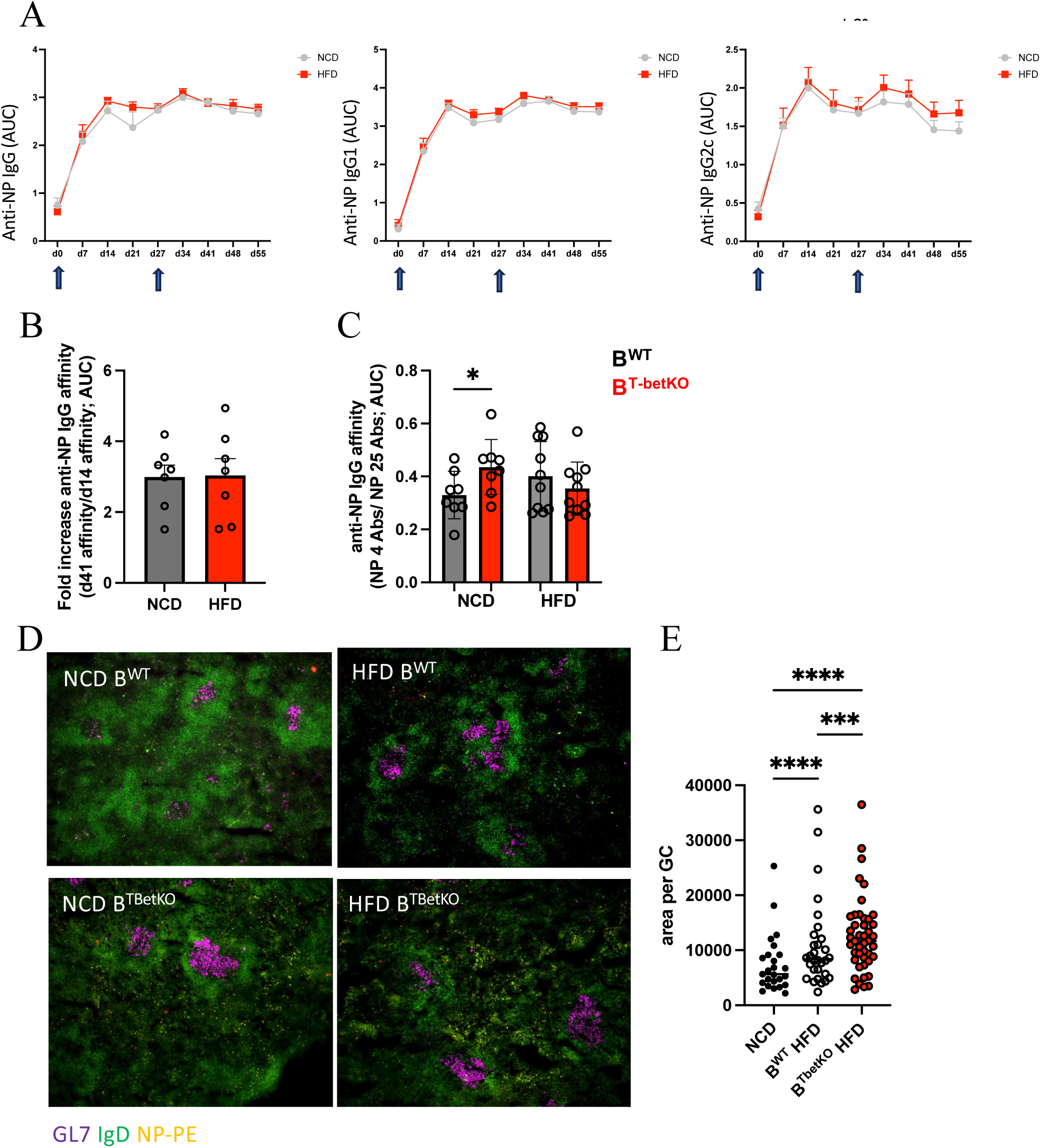
Lean and obese WT B6 mice develop similar antibody titers but B^TbetKO^ mice develop increased antibody affinity compared to B^WT^ controls in lean but not obese mice following NP-KLH/alum immunization. Serum from mice immunized IP with NP-KLH/alum on day 0 and day 27 was collected weekly from WT B6 mice fed a NCD or HFD and was tested by ELISA for anti-NP IgG, IgG1, and IgG2c antibody titers (**A**) as well as fold increase in affinity of IgG anti-NP serum antibody between day 14 to day 41 (**B**). NCD and HFD B^TbetKO^ and B^WT^ mice were examined 14 days after NP-KLH/alum immunization for affinity of IgG NP-specific serum IgG (**C**), immunofluorescent detection of splenic CD19+ (green) GL7+ (magenta) GCs and NP-specific B cells (yellow) [representative images in (**D**)], as well as quantification of area of GCs from immunofluorescent images as in D (**E**). One experiment (**A,B**), or two independent experiments (**C,D,E**). Symbols represent individual GCs (**E**). n=5-7 female mice/grp. Bars indicate mean ± SEM; [Unpaired t test (**C**); Two way ANOVA (**E**)] *p < 0.05, *** p<0.001; ****p < 0.0001.

## DISCUSSION

B cells are a central immune player in obesity and in the progression of its many comorbid disorders. Through the production of antibodies, cytokines, antigen presentation, and the likely modulation of T cells and macrophages, B cells promote the chronic inflammation driving the development of insulin resistance and other metabolic dysfunctions in obesity ^70^. Additionally, as highlighted by the recent COVID-19 pandemic ^2,71,72^, obesity profoundly impacts the effective generation of adaptive immune responses to infection and vaccination ^4,73–75^. Given the increasing prevalence of obesity in adults and children worldwide ^76,77^, understanding the factors that facilitate the collaboration between T cells and B cells in the setting of chronic inflammation will be critical for optimizing vaccination responses and immune defense in patients with obesity or other chronic inflammatory conditions.

Our previous studies identified an association between obesity and the upregulation of the transcription factor T-bet in human and murine B cells, with a resulting expansion of CD11c+ T-bet+ B cells in spleen. We now identified a similar expansion in adipose tissue and liver but not omentum or mesenteric fat following HFD feeding. We had previously discovered that T-bet+ B cells promoted inflammation and metabolic dysfunction in a HFD-induced mouse model of obesity ^53^. Here, we extend those findings to show that eliminating T-bet+ from B cells allows for an increase in obesity-driven autoimmune responses and vaccine-elicited GC B cell responses.

In this study, we observed a significant accumulation of circulating IgG immune complexes (ICs) in the serum of obese mice. This accumulation correlated with the total animal weight, particularly in female mice. Additionally, while both male and female obese mice accumulated IgG1 ICs, only female mice had a significant accumulation of ICs containing the antibody isotype produced predominantly by Tbet+ CD11c+ B cells: IgG2c. During obesity, increased adipocyte death leads to release of cell-free nucleic acids and other self-antigens, elevation of self-antigen-specific autoantibodies in the adipose tissue and in circulation ^78–80^ ^78,81^ ^54,80,82^, and a perpetuation of autoimmune adaptive responses in secondary lymphoid organs^83^. ICs isolated from obese mice stimulate inflammatory cytokines such as TNFa and IL-6 responses from leukocyte populations isolated from adipose tissue, which could be an early or initiating driver of inflammation during obesity. The class-switched IgG1 and IgG2c in ICs from obese mice recognize autoantigens DNA and RNA, suggesting these IC Abs may result from activation of autoreactive B cells which received help from a T cell population. This led us to investigate both the increased level of B cell activation and parallel changes in helper T cell subsets during obesity.

During the course of HFD feeding in mice, their splenic B cell compartment enlarged, and analysis across splenic B cell populations revealed much of that expansion was attributable to the GC and CD11c+Tbet+ B cell populations. T-bet+ B cells in naive lean mice have a higher percentage of class-switched cells compared to other B cells, but even more T-bet+ B cells become class-switched during the HFD, consistent with an antigen-driven, T-cell supported response. We observed the highest B cell expression of CD86 and MHC-II in the T-bet+ B cell population, suggesting these cells are well suited for interactions with helper T cells. Furthermore, Tbet+ B cells from obese animals expressed a high level of scavenger receptor CD36, maintained higher levels of neutral intracellular lipids, and survived high lipid concentrations better than their Tbet-counterparts. Recent studies showed that lipid uptake by CD36 on B cells is critical for fueling mitochondrial OXPHOS and autoreactive B cell differentiation during SLE ^84^, suggesting CD36 on T-bet+ B cells could play a key role in supporting downstream B cell metabolic changes and preferential differentiation of T-bet+ B cells during obesity. Additionally, CD36 could serve as a source of antigen uptake for CD11c+Tbet+ B cells, which is the subject of ongoing studies. Together, these results support the idea that CD36 contributes to antigen-driven activation of T-bet+ CD11c+ B cells and helps to position them to present antigens for engagement with T cells for help.

To address this question, we quantified T helper cells during obesity and found that HFD-induced increases in PD-1^HI^ CXCR5^pos^ Bcl-6+ CD4+ T_FH_ and PD-1^HI^ CXCR5^neg^ Bcl-6-CD4+ T_PH_ cells correlated strongly with elevated T-bet+ B cell frequencies in the spleen and liver, respectively. We found no increase in iNKT_FH_/_PH_ cells in the liver or spleen after 12 weeks of HFD; but iNKT cells in both tissues did express high levels of T-bet and produced abundant amounts of IFNγ^12^, confirming our previously published description of their early contribution to T-bet+ B cell formation. Equivalently high expression of the inducible T cell costimulator (ICOS) molecule in both T_FH_ and T_PH_ cells lends further credence to the B-helper function of these populations. In addition, both subsets of PD-1^HI^ CD4+ T cells were simultaneous producers of IFNγ and IL-21, cytokines needed for T-bet+ B cell generation ^25,36–38^. T-bet was coincident with Bcl-6 expression in T_FH_ cells and expressed in T_PH_ cells which were also the predominant producers of cytokines, suggesting both Tbet+ helper populations would be considered T_H1_ subsets. Such T_FH1_ and T_PH_ populations of cells were previously demonstrated to drive T-bet+ B cell expansion in an *Erlichia muris* model of infection ^38^, and both cell types have been shown to promote T-bet expression in memory B cells in vitro ^25,48^. Our findings therefore suggest that obesity results in expanded T cell helper pools biased towards the generation of T-bet+ B cells. Given this data, plus the preponderance of evidence for the dependence of T-bet+ B cell generation on T cell-help ^25,38,39,41^, it follows that T_FH1_ and T_PH_ cells are likely to support T-bet+ B cell expansion during obesity.

Furthermore, the differentiation and maintenance of T_FH_ cells requires reciprocal interactions with B cells, and T-bet+ B cells appear well-suited to partner with T_FH1_ and T_PH_ cells, specifically. T-bet+ B cells localize to the T/B border within secondary lymphoid organs ^85^ and accumulate within inflamed peripheral tissues; geography which may greatly foster interactions with T_FH1_ and T_PH_ cells ^45,86^. T-bet+ B cells are also widely considered to be antigen-experienced cells ^14^, and both T_FH1_ and T_PH_ cells preferentially help antigen-experienced B cells ^87,88^. This interaction may mirror similar dynamics observed in a cell-intrinsic model of lupus, wherein excessive generation of CD11c+ T-bet+ B cells was linked to aberrantly increased T_FH_ cell differentiation and impaired antigen-specific GC responses ^89^. In testing for reciprocal interactions between T-bet+ B cells and T_FH_/T_PH_ cells, we found that HFD B^TbetKO^ mice had greater numbers of splenic PD-1+ ICOS+ CD4+ T cells. This increase was primarily observed in the splenic T_PH_ compartment, but the levels of T_FH_ cells in both HFD B^TbetKO^ and LC mice were still similar to HFD WT mice. We reasoned that T-bet+ B cells may therefore modulate T_FH_/T_PH_ activity and/or localization, prompting T_FH_ localization away from the follicle to support extrafollicular T-bet+ B cell responses and influencing the egress of T_PH_ cells from secondary lymphoid organs through a common mechanism.

Both T_FH_ and T_PH_ cells in HFD-fed mice expressed high levels of CXCR3, a migratory marker that is associated with trafficking to sites of peripheral inflammation ^43,90^. T-bet+ CD11c+ B cells are important B cell-derived sources of CXCL10 and produce more of this CXCR3-ligand than do conventional B cells^12^. Recently a population of CXCL9/10-producing T-bet+ B cells associating with highly activated CXCR3+ T_H1_ cells was identified in the synovial fluid of patients with rheumatoid arthritis ^91^. Coculture with T_H1_ cells induced CXCL9/10 expression in human B cells from RA patients, and B-cell derived CXCL9/10 in turn significantly induced CD4+ T cell migration. To examine whether CXCR3 signaling was similarly critical for mediating interactions between T-bet+ B cells and T_FH_/T_PH_ cells during obesity, we blocked CXCR3 signaling in HFD wild-type mice. Interestingly, CXCR3 blockade phenocopied HFD B^TbetKO^ mice: the number of splenic, but not hepatic, PD-1+ ICOS+ CD4+ T cells were increased, specifically Tbet+ T_FH1_ and T_PH_ cells. Therefore, we suggest that T-bet+ B cells promote T_PH_ cell egress from the spleen via CXCR3-dependent signaling. CXCR3-blockade also increased GC B cells and class-switched B cells, suggesting more efficient intrafollicular T cell-derived help. CXCR3 was found to play an opposite role during flu infection ^92^, but this dichotomy may specifically highlight different roles for germinal center T-B cell interactions during acute flu infection vs extrafollicular T-B interactions during the chronic inflammation of obesity. These data suggest that during the chronic inflammation of obesity, CXCR3-dependent signaling promotes likely extrafollicular interactions between T-bet+ B cells and T_PH_ cells, and interference via antibody-mediated blockade or absence of CXCL10-producing T-bet+ B cells reroutes T_H_ cells towards providing intrafollicular GC help.

We next examined whether the enhanced GC activity observed in the obese B^TbetKO^ mice mimicked the abnormal B cell responses which arise in certain infectious and autoimmune contexts that develop disrupted germinal centers and/or prolonged EF reactions ^10,11,93^. This skewed response initially favors rapid lower-affinity responses at the expense of longer lived higher affinity responses producing impaired immunity against infections and autoimmunity^10,11^. In obese mice, we see a significant effect of Tbet deficiency in B cells (B^TbetKO^), which produces increases in class-switched B cells, GC B cells, CD138+ IgD-plasma blasts plus accompanying reductions in CD11c+ (extrafollicular) B cells only in mice on a HFD, not NCD controls. Nonetheless, similar but much lower magnitude differences may be possible to observe in NCD mice (since they also possess Tbet+ B cells, but at a lower frequency) if we tested a much larger number of mice. This may also be more evident in older NCD mice, since the background number of Tbet+ B cells is higher in NCD-fed older mice compared to NCD-fed younger mice.

We also considered whether this effect was generalized to vaccine antigens administered during the chronic inflammation of obesity. After immunizing obese B^TbetKO^ and B^WT^ mice with the T-dependent antigen NP-KLH plus alum, we saw increases in NP+ B cells, NP+ class switched B cells, and NP+ GC B cells in B^TbetKO^ mice compared to B^WT^ mice, when fede both a NCD and HFD. Importantly, all mice on NCD and HFD in Figures 7 and 8 were immunized with NP-KLH plus alum adjuvant, creating an inflammatory environment in both sets of mice. This expands all B cells in general as well as Tbet+ B cells, and the data suggests this expansion is enough in even the NCD mice for the loss of Tbet in B cells to lead to a detectable effect on GCs. The NP-KLH vaccination system does not permit antibody neutralization testing to confirm increased effectiveness from the enlarged GCs, but the role of obesity-expanded Tbet+CD11c+ B cell effects on antibody effector mechanisms will benefit from further testing in the context of a murine infectious disease model where antibody neutralization capability can be directly assessed. In summary, mice who lack T-bet expression in B cells are able to mount expanded antigen-specific GC B cell responses to an immunization with a T-dependent antigen, and this effect can be exacerbated by obesity. By extension, obesity-expanded T-bet+ B cell inhibition of autoantigen-reactive and foreign antigen-reactive B cell GC responses may be restored by eliminating T-bet selectively from the B cell population. This warrants further comparative study and expands the relevance of these results to broad circumstances of chronic inflammation beyond obesity. This includes aging and autoimmunity, which are also characterized by expanded Tbet+ B cells, where patients exhibit the same reduced immune protection observed during obesity.

These results fit with a recently proposed model of B cell responses during obesity which reconciles the presence of a proinflammatory and hyperactivated B cell phenotype together with the impaired generation of protective humoral immunity against pathogens^10^. Previous understanding has been that in lean individuals B cell responses are typically characterized by early and short-lived EF reactions preceding germinal center responses which then generate long-lasting and higher-affinity antibodies and memory B cells. The new model proposes that in obese individuals, B cell responses favor EF reactions with increased magnitude and duration which will impair optimal GC responses, leading to the reduction of protective long-lasting antibodies and memory B cells ^10^. Our results support this model, where expression of T-bet expands a highly activated extrafollicular population representing a substantial portion of the IgG-expressing B cell repertoire. Consistent with this idea, genetically engineered elimination of T-bet expression in B cells skewed subsequent immune responses to favor GC reactions, which augments the ability of B cells to class-switch and differentiate into antibody-secreting cells. Compelling and detailed mechanistic support for this interpretation has been provided by the Shlomchik laboratory’s recent publication demonstrating that during SLE, B cellsdirectly inhibit GCs via a feedback loop of B and T cell-derived IFNγ and IL-12 ^69^. Our results are consistent with a similar mechanism operating during obesity. The role of T-bet+ B cells in obesity may have broad implications since these cells are also expanded in elderly and autoimmune individuals, where they are enriched for autoreactive specificities ^13,15,20,24,37,94,95^. The human equivalent of T-bet+ B cells (double negative 2 B cells, IgD-CD27-CXCR5-CD11c+) also produce autoantibodies during obesity ^35,79,80,96^.

In summary, these results support the hypothesis that B cell activation in obesity results in the expansion of T-bet+ B cells which produce autoantibodies, generate inflammatory ICs, and support T_FH_/T_PH_ differentiation. Genetically eliminating T-bet in B cells reduces the levels of IgG2c ICs and autoantibodies, suggesting they are CD11c+ Tbet+ B cell derived. Mice lacking T-bet expression in B cells develop expanded GC reactions, and increased T_FH_/T_PH_ cells during obesity and following vaccination. However, lack of Tbet in B cells only led to increased antibody affinity in response to vaccination in lean mice so questions remain. It will be useful to evaluate clinically relevant infections to determine if the antibodies produced in the absence of CD11c+Tbet+ B cells in lean or obese mice are more effective. Finally, these findings suggest a possible immunologic mechanism for patients with expanded T-bet+ B cell populations in spleen and adipose tissue who produce less effective humoral immune responses to vaccines and suffer from worse clinical outcomes during infections.

### Limitations of the Study

Male mice consistently develop obesity when fed a HFD and are commonly used for obesity studies; however, this limits our understanding of obesity to half the population. Our previous studies identified increases in Tbet+ B cells in female human adipose samples. While we observed similar changes in Tbet+ B cells in male and female mice, the magnitude of the B cell phenotype was more evident in females. To match our previous studies, we started here with a careful analysis of changes in immune populations across various secondary and tertiary immune tissues from female mice fed a HFD starting at 8 or 16 weeks of age. Starting the HFD in female mice at 16 weeks of age optimized consistent weight gain in this model, so this timeline was used in all remaining studies. Future studies could determine if male mice fed a HFD would exhibit the same immune cell changes observed here in female mice.

This study identifies the unique expansion of Tbet+ B cells as well as T_FH_ and T_PH_ cells, but not iNKT_FH_ or iNKT_PH_ cells, in spleen and liver, respectively, during obesity induced by HFD. The study does not attempt to isolate or phenotype similar populations in NCD-fed mice because the same cells are extremely rare (i.e. not expanded) and perhaps more artifactual in NCD-fed mice where there is no driving inflammation. It was beyond the scope of the current study to compare T_FH_ and T_PH_ cells in the chronic inflammatory state of obesity with T_FH_ or T_PH_ cells from an acute infection, but this will be a very interesting future comparison.

## ACKNOWLEDGMENTS

We would like to thank Sebastian Montagnino, Catherine Davis, and Dr. Yue Li of the UT Health Flow cytometry Core for exceptional technical support. We would also like to thank the UT Health San Antonio Laboratory Animal Resources and vivarium staff for their essential services, as well as the US NIH Tetramer Core for CD1d-PBS57 tetramers. We are also grateful to Dr. Nu Zhang for providing mice, as well as Drs. Gary Winslow, Laurence Morel (UTHSCSA), Lily Q. Dong (UTHSCSA), Mark J. Shlomchik (UPitt), and Mikael Karlsson (Karolinksa Institutet) for their fruitful discussions and kind assistance.

## FUNDING

The Flow Cytometry Shared Resource at UT Health San Antonio is supported by the National Cancer Institute (P30CA054174) through the Mays Cancer Center, the Cancer Prevention and Research Institute of Texas (CPRIT) (RP210126), the National Institutes of Health (S10OD030432), and the Office of the Vice President for Research at UT Health San Antonio. This work was also supported by critical funding from the National Institutes of Health: grant R01 AI32789-01A1 (EAL), K12 GM111726 (BTE).

## AUTHOR CONTRIBUTIONS

CV, BTE, and EAL conceived the project, CV, BTE, and EAL planned the approach, while CV, BTE, EC, MDL, NL, and EAD performed all experiments. Funding was acquired by BTE and EAL. Project administration as well as most mentoring and supervision was provided by EAL. CV mentored and supervised NL; EC mentored and supervised MDL. The manuscript was written and edited by CV, BTE, and EAL.

## STAR METHODS

### Diet-induced obesity model

Mice were fed with autoclaved standard pellet chow (NCD) or 60 kcal% fat chow (HFD, Research Diets) and water *ad libitum*. Mice for the DIO model were initiated on NCD or HFD at either 6-10 or 14-16 weeks of age and maintained on the diet for 11-14 weeks. Bedding was mixed 1 week prior to initiation of NCD/HFD among different cages. Mouse weight and food intake were recorded bi-weekly.

### Mice

C57BL/6J wild-type mice (#000664) and B6.129P2(C)-Cd19tm1(cre)Cgn/J (CD19^Cre^; #006785) were obtained from The Jackson Laboratory. B6.129-Tbx21tm2Srnr/J mice (*Tbx21*^flox^ Jax#022741) were obtained from Dr. Nu Zhang (UT Health San Antonio). *CD19*^Cre/+^ and *Tbx21*^flox/flox^ mice were intercrossed to generate *Tbx21*^flox/flox^ *CD19*^Cre/Cre^ mice. *Tbx21*^flox/flox^ *CD19*^Cre/Cre^, *Tbx21*^flox/flox^, and WT mice were intercrossed to generate *CD19*^Cre/+^ (T-bet+ B LC; B^WT^) and *Tbx21*^flox/flox^ CD19^Cre/+^ (T-bet+ B KO; B^TbetKO^) mice. All mice were housed and bred under specific pathogen-free conditions at the animal facility of UT Health (UTHSCSA LAR). Age- and sex-matched female and male mice at 18-25 weeks of age were used in experiments in **Fig 2A**. All other experiments only included female age-matched mice at 16-30 weeks of age (for *in vivo* experiments) and 8-12 weeks of age (for *in vitro* experiments). Mice were fed a high-fat diet (HFD), in which 60% of kcal% was fat (Research Diets Inc.). Feeding was allowed *ad libitum,* and mouse weights and food intakes were monitored twice weekly. 1 week prior to assignment to experimental and control groups and start of diets, bedding was exchanged between all cages. All experiments were performed under UT Health Animal Care and Use Committee-approved protocols.

### Murine tissue processing

Mouse spleens were processed by mechanical disruption using 100 μm metal strainers to generate single-cell suspensions. Mouse peri-gonadal white adipose tissue (WAT), omental tissue, and mesenteric adipose tissue (MAT) was dissected and minced with fine scissors before digestion with type-IV collagenase (0.1 mg/mL) in RPMI for 30 min at 37°C on an orbital shaker. For both tissues, red blood cells were lysed using incubation with ACK buffer (ThermoFisher Scientific). Single-cell solutions were filtered through 100 um (WAT) or 40 um (spleen) cell strainers and cells were collected by centrifugation. Murine livers were first perfused with 5 mL of 1X phosphate buffered saline (PBS) (Gibco^TM^) via injection through the portal vein. Liver tissue was passed through a 100 μm metal strainer then suspended in a 40% Percoll (Cytiva) solution, underlaid with 60% Percoll/RPMI, and centrifuged. Lymphocytes at the interface of the Percoll gradient layers were collected with a transfer pipette and washed with RPMI. For all tissues, cell yields and viability were measured by trypan blue staining using a hemocytometer.

### Injections

NP-KLH and Alum were emulsified before intraperitoneal (IP) administration by diluting stock NP-KLH solutions to 1 mg/mL in PBS, then adding an equal volume of Alum dropwise while stirring for 30 minutes. 200uL injection per mouse contained 100ug NP-KLH in Alum. C57BL/6J WT female mice were injected IP with 0.25 mg of anti-mouse CXCR3 (CD183) and anti-mouse CXCL9 (MIG) or hamster IgG (BioXCell) every other day for 14 days after completing 12 weeks on HFD.

### In vitro splenocyte and SVF assays

Splenocytes in single-cell suspensions were harvested from WT female mice at 8-12 weeks of age. B cells were isolated from total splenocytes using the EasySep Mouse Pan-B Cell Isolation Kit per the manufacturer’s instructions. 1-2 x 10^6^ B cells were plated in 48- or 96-well flat-bottomed plates and cultured in complete RPMI supplemented with FBS, L-glutamine, HEPES, Pen-Strep and 2-Mercaptoethanol. For free fatty acid (FFA) treatment assays, B cells were stimulated using a cocktail of α-CD40 (1 ug/mL), R848 (10 ug/mL), IL-21 (25 ng/mL) and IFNγ (10 ng/mL) and treated with either BSA-conjugated oleate, palmitate, or free BSA at 25, 100, and 400 uM. The cells were cultured for 3 days prior to flow cytometry analysis. For SVF cultures, cells were added to 96 well plate coated with equal quantity column-purified IgG from NCD or HFD fed mice, cultured for 24hrs, and supernatants collected for ELISA analysis. TNFα and IL-6 ELISA kits (Biolegend) were used per manufacturer instructions.

### *Ex vivo* stimulation and intracellular cytokine staining

Isolated hepatic and splenic leukocytes from HFD female mice were plated in 96-well flat-bottomed plate and cultured at a density of 1.0 x 10^6^ cells/mL at 37°C in RPMI supplemented with 10% FBS, L-glutamine, HEPES, Pen-Strep, and 2-Mercaptoethanol (ThermoFisher Scientific). Cells were stimulated with eBioscience^TM^ Cell Stimulation Cocktail in the presence of monensin and brefeldin A (Biolegend) for four hours before flow cytometry analysis. After fixation/permeabilization, cells were labeled with recombinant mouse IL-21 receptor/human Fc chimera protein (R&DSystems Inc.), followed by R-phycoerythrin-conjugated AffiniPure F(ab’)2 Fragment Goat Anti-Human IgG (Jackson ImmunoResearch), and then BV786-labeled anti-IFN-γ.

### ELISA

Mouse serum was isolated from blood collected via submandibular vein bleed or terminal eye bleed. After clotting at room temperature, blood was centrifuged twice at 10,000 rpm for 6 minutes and serum stored at −80C until analysis. ELISA detecting antibodies included HRP-conjugated anti-mouse IgM, IgG, IgG1, IgG2b, IgG2c, and IgG3 antibodies (Southern Biotech). ABTS or TMB were substrates for the HRP. Human C1q (10 ug/mL), calf thymus DNA (50 ug/mL), human oxLDL (5 ug/mL), mouse liver RNA (50 ug/mL), and NIP-OVA (5ug/mL) were used to coat the wells of the Costar ELISA plates to detect immune complexes, α-DNA, α-oxLDL, α-RNA, and α - NP antibodies, respectively. For affinity ELISAs, wells of ELISA plates were coated with NP^4^-BSA or NP^25^-BSA.

### Flow cytometry

Once single-cell suspensions were obtained from mouse tissues *ex vivo* or from *in vitro* cultured cells, samples were treated with purified anti-mouse CD16/32 antibody (eBioscience) for 10-15 minutes to block non-specific binding. Cells were also stained with viability dyes (Zombie NIR Fixable Viability Kit (Biolegend) or LIVE/DEAD^TM^ Fixable Blue Dead Cell Stain Kit (Invitrogen)) for 10 min at room temperature. Cells were then washed and stained for extracellular markers using indicated antibodies for 30 minutes, in the dark, at 4°C. To readily identify CXCR5+ cells, samples were first stained with Biotin Rat Anti-Mouse CD185 (CXCR5) antibody along with other extracellular makers. Following washing, samples were then stained with Streptavidin-APC/Cy7 for 30 min at 4°C. For iNKT cell identification, cells were stained for 25 min using PBS57-loaded CD1d tetramers at room temperature on an orbital shaker, followed by two additional wash steps. Samples were then washed and stained for 30 minutes at 4C with the indicated antibodies to label extracellular markers. For staining of intracellular markers, samples were fixed and permeabilized using the eBioscience Fixation & Permeabilization Buffer Set according to the manufacturer’s instructions. Samples were then washed and stained for 30 minutes at 4C with the indicated antibodies to label intracellular markers. Finally, samples were washed, filtered (WAT), resuspended in 2% FBS in PBS, analyzed, and gated on a BD FACSCELESTA^TM^ or Cytek Aurora flow cytometer, as depicted (**Fig. S8)**. Doublet exclusion, debris exclusion, and live-cell discrimination were performed on all samples prior to additional flow cytometry gating and analysis. Data were analyzed using BD FACSDiva^TM^, SpectroFlo (Cytek Biosciences), and FlowJo 10.7.1 software.

### Antibodies used in this study

The following anti-mouse antibodies were used in this study: F4/80 Brilliant Violet^TM^ (BV)421 (Bm8), CD45 BV786 (30-F11), CD45R/B220 BV510 or PerCp-Cy5.5 (RA3-6B2), T-bet phycoerythrin (PE)-Cy7 (4B10), TRCβ BV650, BV510, or BV605 (H57-597), CD183/CXCR3 BV650 (CXCR3-173) were obtained from BioLegend. F4/80 Brilliant Ultra Violet^TM^ (BUV)661 (T45-2342), CD19 BV786, BV650, or BUV395 (1D3), CD11c PE or allophycocyanin (APC) (HL3), or CD4 fluorescein isothiocyanate (FITC) (RM4-5), CD279/PD-1 BUV737 (RMP1-30), IFNγ BV786 (XMG1.2) and Bcl-6 BV421 (K112-91) were obtained from BD Biosciences. CD25 FITC (7D4) was purchased from SouthernBiotech. CD4 eFluor^TM^ 615 (RM4-5) and CD278/ICOS PE (7E.17G9) was purchased from eBioscience. CXCR5 staining was performed by using BD Pharmingen^TM^ Biotin Rat Anti-Mouse CD185 (CXCR5) (2G8) followed by Streptavidin APC-Cy7 from BD Biosciences. iNKT cells were identified using tetramers of PBS57-loaded CD1d tetramer (kindly provided by the NIH tetramer core facility) conjugated to APC, PE, or BV421. Intracellular cytokine staining for IL-21 was performed by using recombinant mouse IL-21 receptor/human Fc chimera protein (R&DSystems Inc.), followed by R-phycoerythrin-conjugated AffiniPure F(ab’)2 Fragment Goat Anti-Human IgG (Jackson ImmunoResearch Laboratories).

### Immunofluorescent staining

Kidneys were collected from wild-type NCD and HFD mice (**Fig. 2**) or spleens from CD19^Cre^ B^WT^ and B^TbetKO^ mice (**Fig. 8**) and immediately frozen in OCT compound. Frozen tissue samples were sliced into 6 um sections with a Cryostat at −20C. Sections were fixed using cold histological acetone and blocked for 1 hour using 5% Rabbit Serum or 5% FBS in PBS. FITC-conjugated anti-mouse IgG was diluted in Antibody Dilution Buffer (1:200) and used to measure IgG deposition in the kidney glomeruli. Anti-IgD (Alexa488), GL7 (Alexa647), and NP-PE were used to label antigen-specific B cells and GCs in spleen sections. ProLong Gold with NucBlue Stain was used to mount slides and to stain for kidney cell nuclei. Prolong Glass Antifade and DAPI stain were used to mount slides and stain for spleen nuclei. Kidney sections were analyzed and imaged by immunofluorescence using a Zeiss Microscope. Spleen images were captured and stitched using the Keyence microscope.

## QUANTIFICATION AND STATISTICAL ANALYSIS

Statistical analyses were conducted by GraphPad Prism using one- or two-tailed non-parametric Student’s t test, 1-way ANOVA, 2-way ANOVA, or simple linear regression as indicated in figure legends with p values of less than 0.05 considered significant and marked by asterisk(s) (\**p* < 0.05, \*\**p* < 0.01, \*\*\**p* < 0.001, and \*\*\*\**p* < 0.0001.). Bar heights and symbol centers indicated group means. Dots represent individual values. Error bars indicate S.E.M.

## Notes

### Competing Interest Statement

The authors have declared no competing interest.

### Summary of Updates

Minor grammatical edits, updated incorrect figure reference in text, added organ-specific labels to sub-panels in Fig 5, moved last results paragraph to discussion

