## Supplemental Figures for "CD11c+ Tbet+ B cells constrain obesity- and vaccination-induced germinal center B cells and T helper cells": 9.1.2025 Cell Reports SUPP FIGURES.pptx

### Slide 1
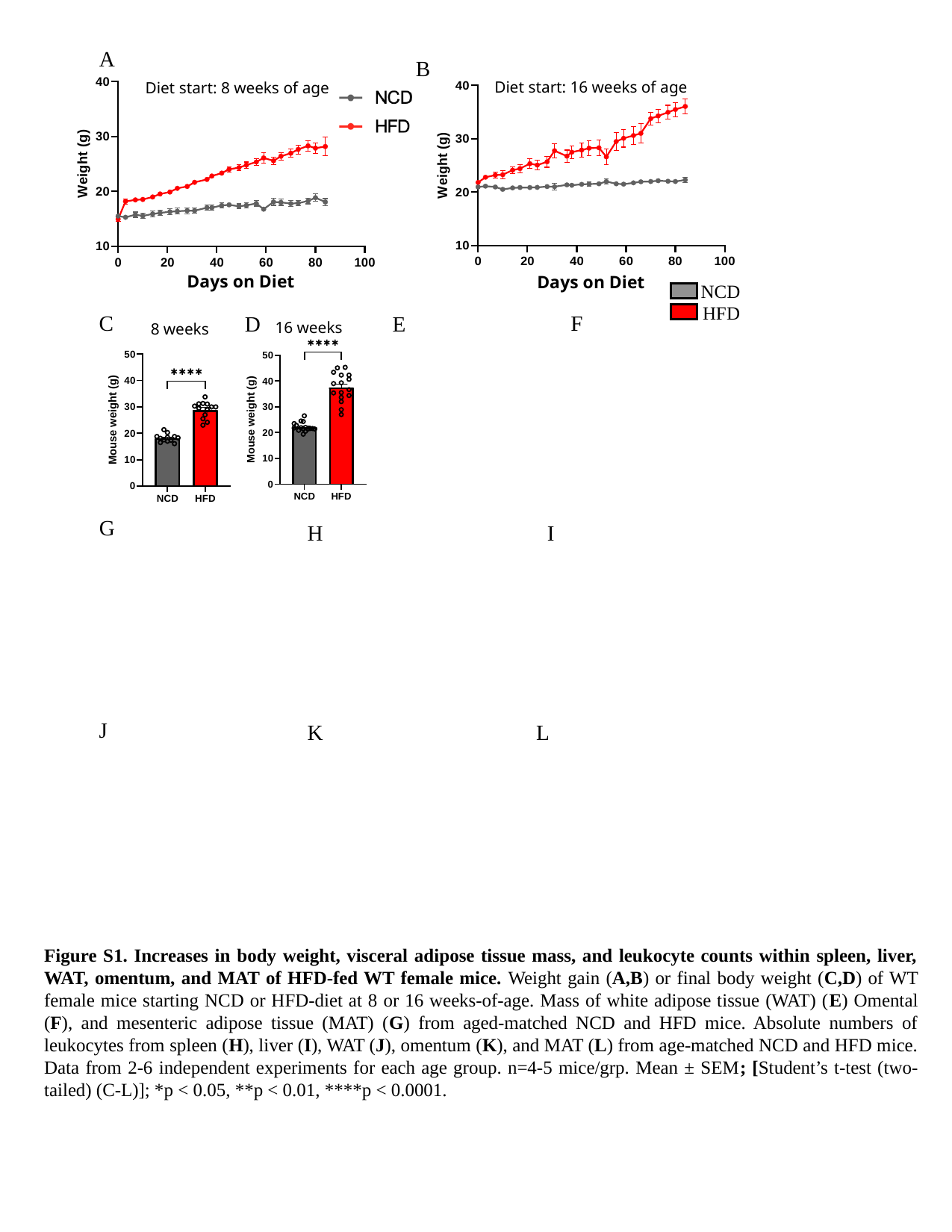

A
B
Diet start: 16 weeks of age
Diet start: 8 weeks of age
Days on Diet
Days on Diet
NCD
HFD
C
F
D
E
16 weeks
8 weeks
G
H
I
J
K
L
Figure S1. Increases in body weight, visceral adipose tissue mass, and leukocyte counts within spleen, liver, WAT, omentum, and MAT of HFD-fed WT female mice. Weight gain (A,B) or final body weight (C,D) of WT female mice starting NCD or HFD-diet at 8 or 16 weeks-of-age. Mass of white adipose tissue (WAT) (E) Omental (F), and mesenteric adipose tissue (MAT) (G) from aged-matched NCD and HFD mice. Absolute numbers of leukocytes from spleen (H), liver (I), WAT (J), omentum (K), and MAT (L) from age-matched NCD and HFD mice. Data from 2-6 independent experiments for each age group. n=4-5 mice/grp. Mean ± SEM; [Student’s t-test (two-tailed) (C-L)]; *p < 0.05, **p < 0.01, ****p < 0.0001.

### Slide 2
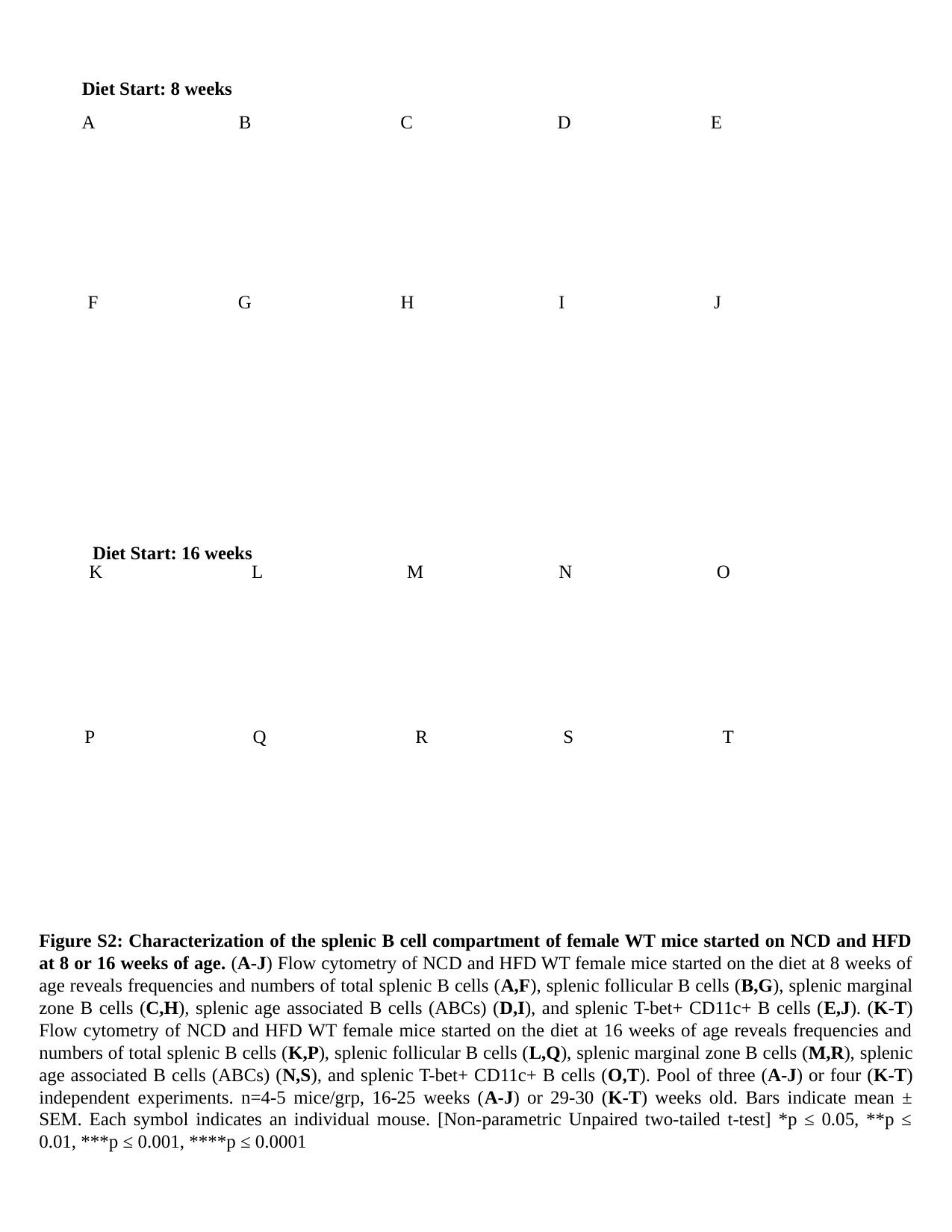

Diet Start: 8 weeks
A B C D E
F G H I J
Diet Start: 16 weeks
K L M N O
P Q R S T
Figure S2: Characterization of the splenic B cell compartment of female WT mice started on NCD and HFD at 8 or 16 weeks of age. (A-J) Flow cytometry of NCD and HFD WT female mice started on the diet at 8 weeks of age reveals frequencies and numbers of total splenic B cells (A,F), splenic follicular B cells (B,G), splenic marginal zone B cells (C,H), splenic age associated B cells (ABCs) (D,I), and splenic T-bet+ CD11c+ B cells (E,J). (K-T) Flow cytometry of NCD and HFD WT female mice started on the diet at 16 weeks of age reveals frequencies and numbers of total splenic B cells (K,P), splenic follicular B cells (L,Q), splenic marginal zone B cells (M,R), splenic age associated B cells (ABCs) (N,S), and splenic T-bet+ CD11c+ B cells (O,T). Pool of three (A-J) or four (K-T) independent experiments. n=4-5 mice/grp, 16-25 weeks (A-J) or 29-30 (K-T) weeks old. Bars indicate mean ± SEM. Each symbol indicates an individual mouse. [Non-parametric Unpaired two-tailed t-test] *p ≤ 0.05, **p ≤ 0.01, ***p ≤ 0.001, ****p ≤ 0.0001

### Slide 3
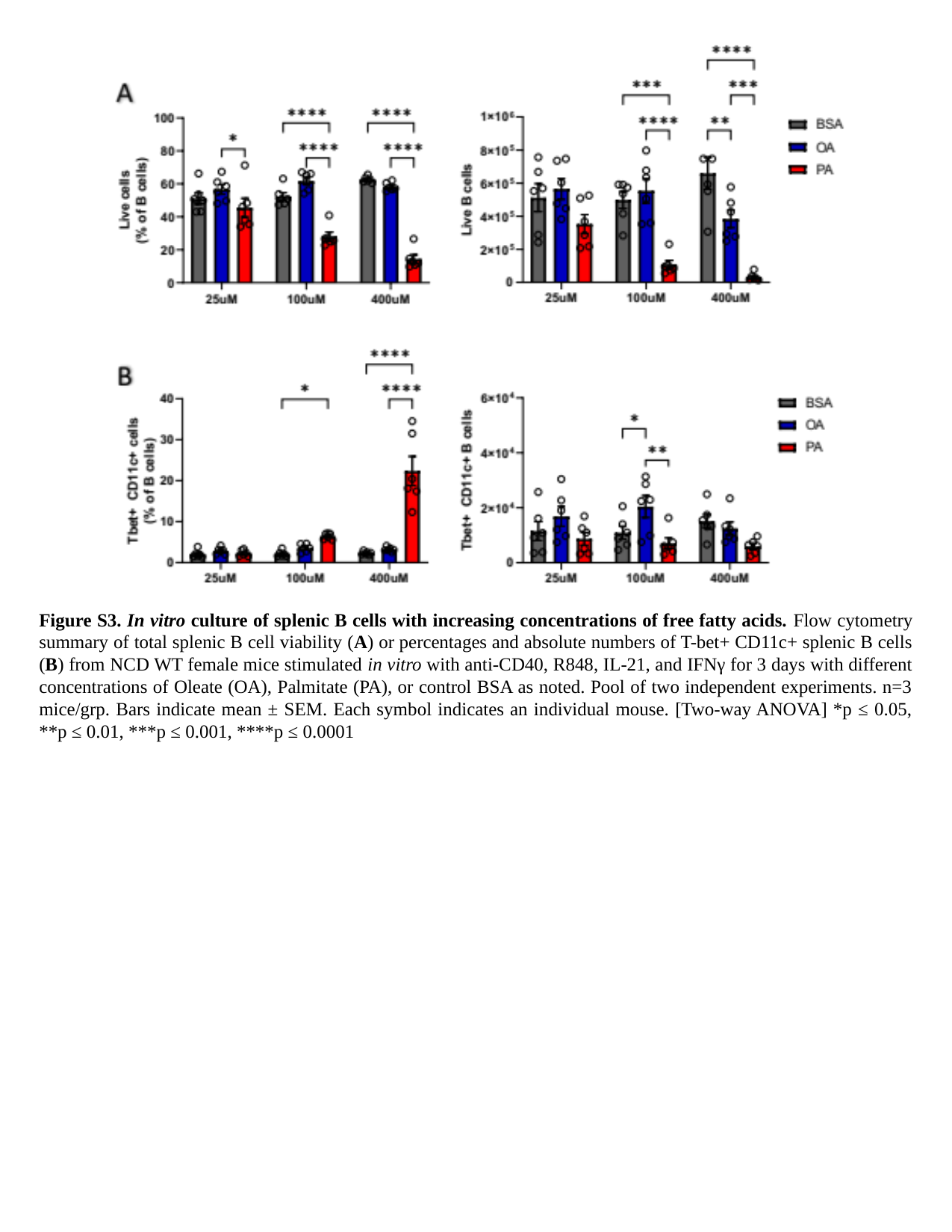

Figure S3. In vitro culture of splenic B cells with increasing concentrations of free fatty acids. Flow cytometry summary of total splenic B cell viability (A) or percentages and absolute numbers of T-bet+ CD11c+ splenic B cells (B) from NCD WT female mice stimulated in vitro with anti-CD40, R848, IL-21, and IFNγ for 3 days with different concentrations of Oleate (OA), Palmitate (PA), or control BSA as noted. Pool of two independent experiments. n=3 mice/grp. Bars indicate mean ± SEM. Each symbol indicates an individual mouse. [Two-way ANOVA] *p ≤ 0.05, **p ≤ 0.01, ***p ≤ 0.001, ****p ≤ 0.0001

### Slide 4
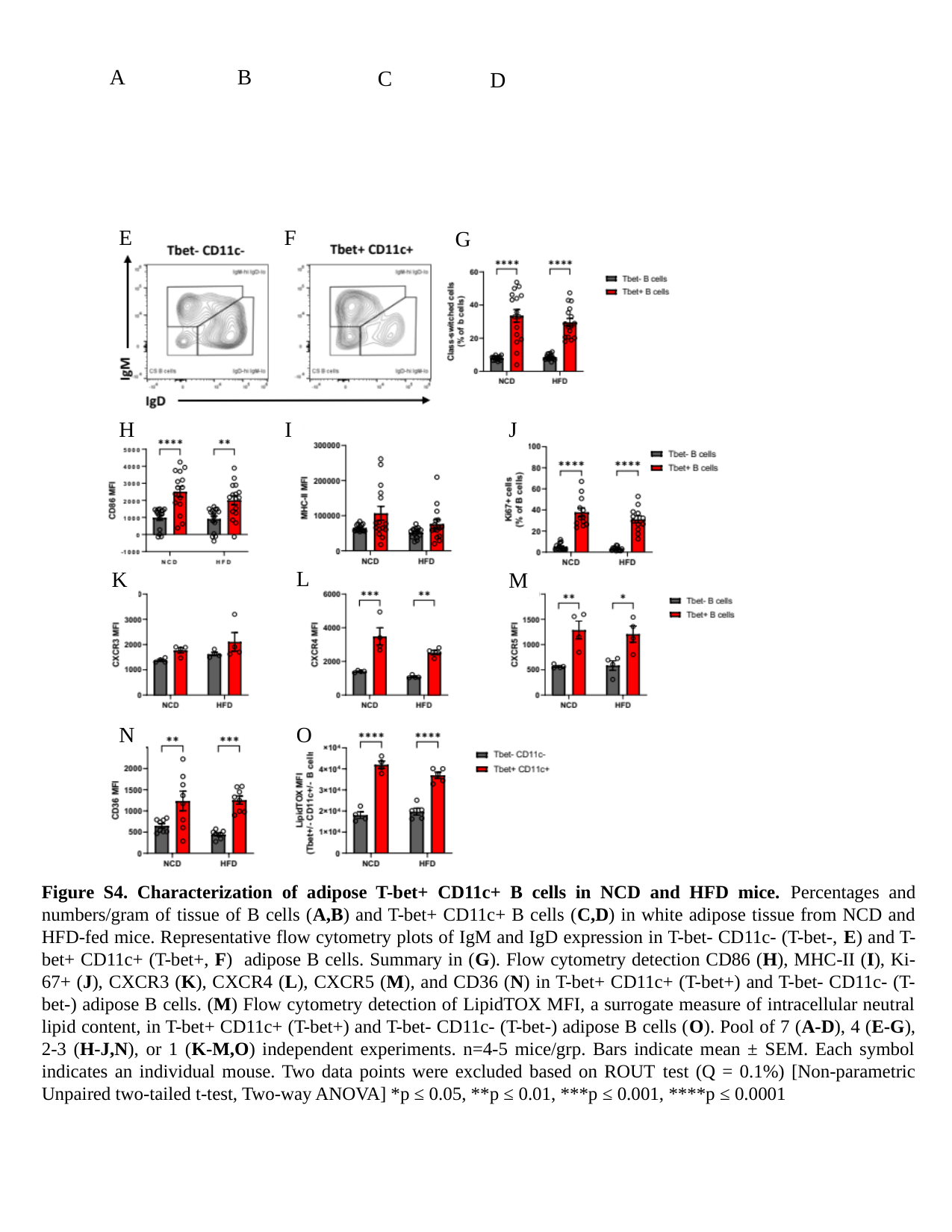

A
B
C
D
E
F
G
H
I
J
L
K
M
O
N
Figure S4. Characterization of adipose T-bet+ CD11c+ B cells in NCD and HFD mice. Percentages and numbers/gram of tissue of B cells (A,B) and T-bet+ CD11c+ B cells (C,D) in white adipose tissue from NCD and HFD-fed mice. Representative flow cytometry plots of IgM and IgD expression in T-bet- CD11c- (T-bet-, E) and T-bet+ CD11c+ (T-bet+, F) adipose B cells. Summary in (G). Flow cytometry detection CD86 (H), MHC-II (I), Ki-67+ (J), CXCR3 (K), CXCR4 (L), CXCR5 (M), and CD36 (N) in T-bet+ CD11c+ (T-bet+) and T-bet- CD11c- (T-bet-) adipose B cells. (M) Flow cytometry detection of LipidTOX MFI, a surrogate measure of intracellular neutral lipid content, in T-bet+ CD11c+ (T-bet+) and T-bet- CD11c- (T-bet-) adipose B cells (O). Pool of 7 (A-D), 4 (E-G), 2-3 (H-J,N), or 1 (K-M,O) independent experiments. n=4-5 mice/grp. Bars indicate mean ± SEM. Each symbol indicates an individual mouse. Two data points were excluded based on ROUT test (Q = 0.1%) [Non-parametric Unpaired two-tailed t-test, Two-way ANOVA] *p ≤ 0.05, **p ≤ 0.01, ***p ≤ 0.001, ****p ≤ 0.0001

### Slide 5
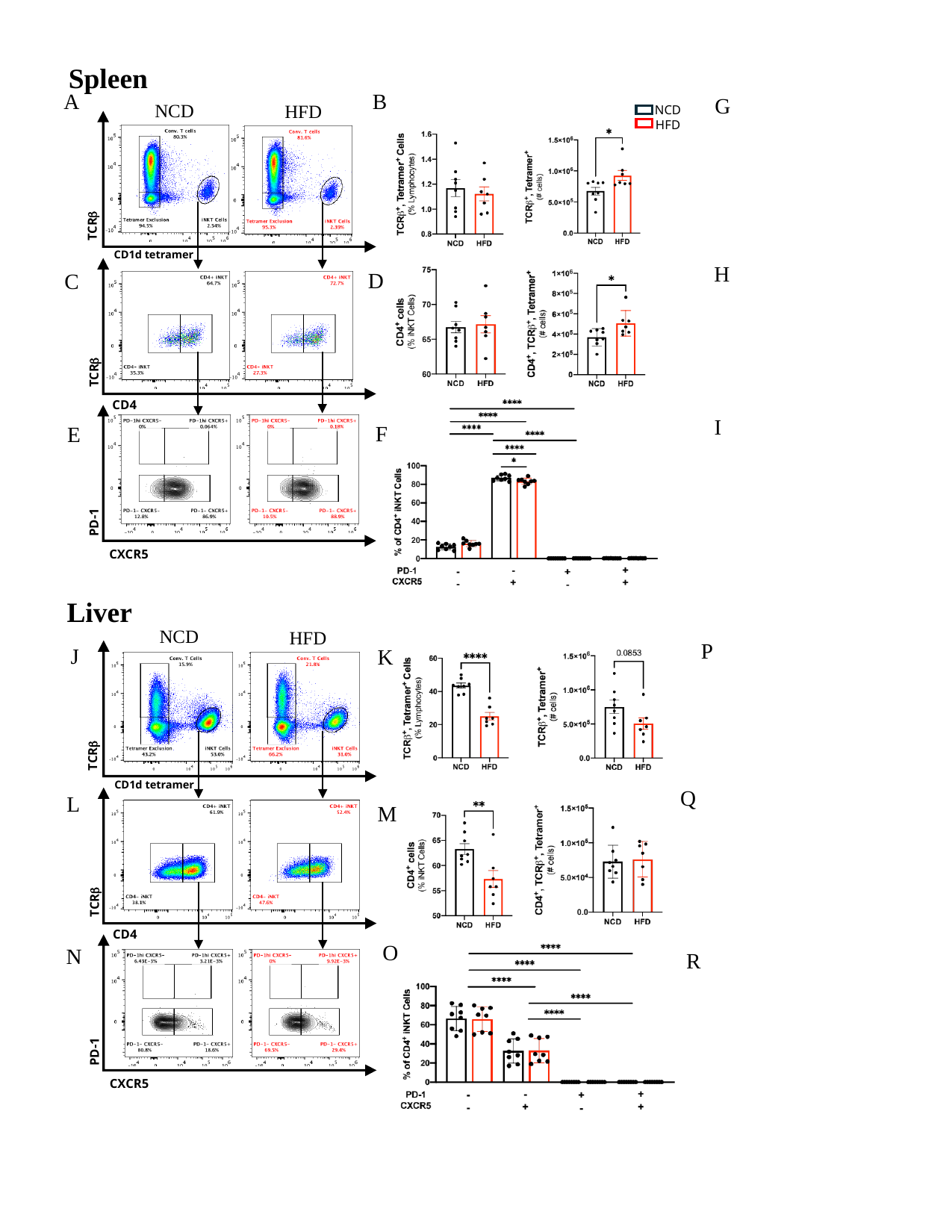

Spleen
A
B
G
NCD
HFD
NCD
HFD
TCRβ
CD1d tetramer
H
D
C
TCRβ
CD4
I
F
E
PD-1
CXCR5
Liver
NCD
HFD
P
J
K
TCRβ
CD1d tetramer
Q
L
M
TCRβ
CD4
O
N
R
PD-1
CXCR5

### Slide 6
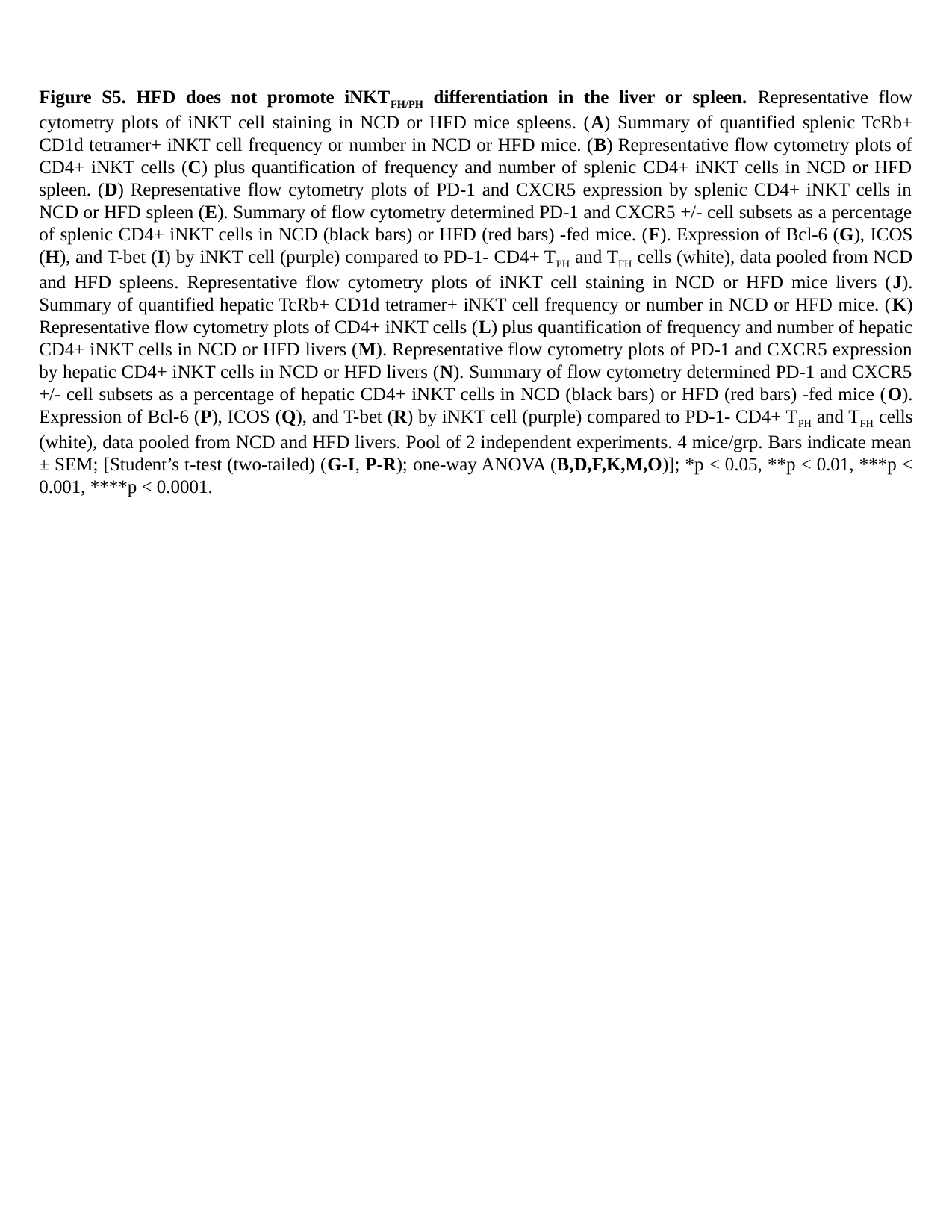

Figure S5. HFD does not promote iNKT_FH/PH_ differentiation in the liver or spleen. Representative flow cytometry plots of iNKT cell staining in NCD or HFD mice spleens. (A) Summary of quantified splenic TcRb+ CD1d tetramer+ iNKT cell frequency or number in NCD or HFD mice. (B) Representative flow cytometry plots of CD4+ iNKT cells (C) plus quantification of frequency and number of splenic CD4+ iNKT cells in NCD or HFD spleen. (D) Representative flow cytometry plots of PD-1 and CXCR5 expression by splenic CD4+ iNKT cells in NCD or HFD spleen (E). Summary of flow cytometry determined PD-1 and CXCR5 +/- cell subsets as a percentage of splenic CD4+ iNKT cells in NCD (black bars) or HFD (red bars) -fed mice. (F). Expression of Bcl-6 (G), ICOS (H), and T-bet (I) by iNKT cell (purple) compared to PD-1- CD4+ T_PH_ and T_FH_ cells (white), data pooled from NCD and HFD spleens. Representative flow cytometry plots of iNKT cell staining in NCD or HFD mice livers (J). Summary of quantified hepatic TcRb+ CD1d tetramer+ iNKT cell frequency or number in NCD or HFD mice. (K) Representative flow cytometry plots of CD4+ iNKT cells (L) plus quantification of frequency and number of hepatic CD4+ iNKT cells in NCD or HFD livers (M). Representative flow cytometry plots of PD-1 and CXCR5 expression by hepatic CD4+ iNKT cells in NCD or HFD livers (N). Summary of flow cytometry determined PD-1 and CXCR5 +/- cell subsets as a percentage of hepatic CD4+ iNKT cells in NCD (black bars) or HFD (red bars) -fed mice (O). Expression of Bcl-6 (P), ICOS (Q), and T-bet (R) by iNKT cell (purple) compared to PD-1- CD4+ T_PH_ and T_FH_ cells (white), data pooled from NCD and HFD livers. Pool of 2 independent experiments. 4 mice/grp. Bars indicate mean ± SEM; [Student’s t-test (two-tailed) (G-I, P-R); one-way ANOVA (B,D,F,K,M,O)]; *p < 0.05, **p < 0.01, ***p < 0.001, ****p < 0.0001.

### Slide 7
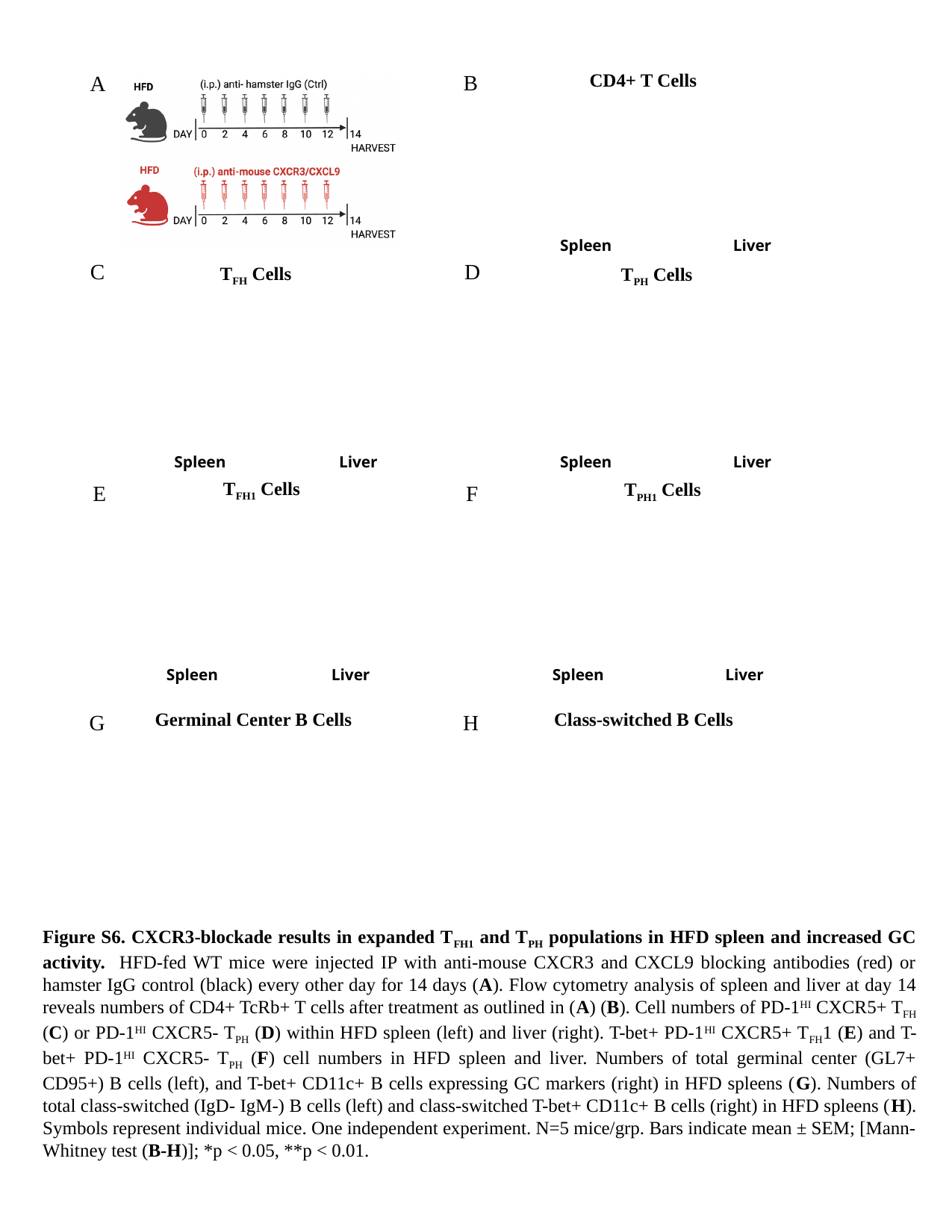

CD4+ T Cells
B
A
Spleen
Liver
D
C
T_FH_ Cells
T_PH_ Cells
Spleen
Liver
Spleen
Liver
T_FH1_ Cells
T_PH1_ Cells
F
E
Spleen
Liver
Spleen
Liver
Germinal Center B Cells
Class-switched B Cells
H
G
Figure S6. CXCR3-blockade results in expanded T_FH1_ and T_PH_ populations in HFD spleen and increased GC activity. HFD-fed WT mice were injected IP with anti-mouse CXCR3 and CXCL9 blocking antibodies (red) or hamster IgG control (black) every other day for 14 days (A). Flow cytometry analysis of spleen and liver at day 14 reveals numbers of CD4+ TcRb+ T cells after treatment as outlined in (A) (B). Cell numbers of PD-1^HI^ CXCR5+ T_FH_ (C) or PD-1^HI^ CXCR5- T_PH_ (D) within HFD spleen (left) and liver (right). T-bet+ PD-1^HI^ CXCR5+ T_FH_1 (E) and T-bet+ PD-1^HI^ CXCR5- T_PH_ (F) cell numbers in HFD spleen and liver. Numbers of total germinal center (GL7+ CD95+) B cells (left), and T-bet+ CD11c+ B cells expressing GC markers (right) in HFD spleens (G). Numbers of total class-switched (IgD- IgM-) B cells (left) and class-switched T-bet+ CD11c+ B cells (right) in HFD spleens (H). Symbols represent individual mice. One independent experiment. N=5 mice/grp. Bars indicate mean ± SEM; [Mann-Whitney test (B-H)]; *p < 0.05, **p < 0.01.

### Slide 8
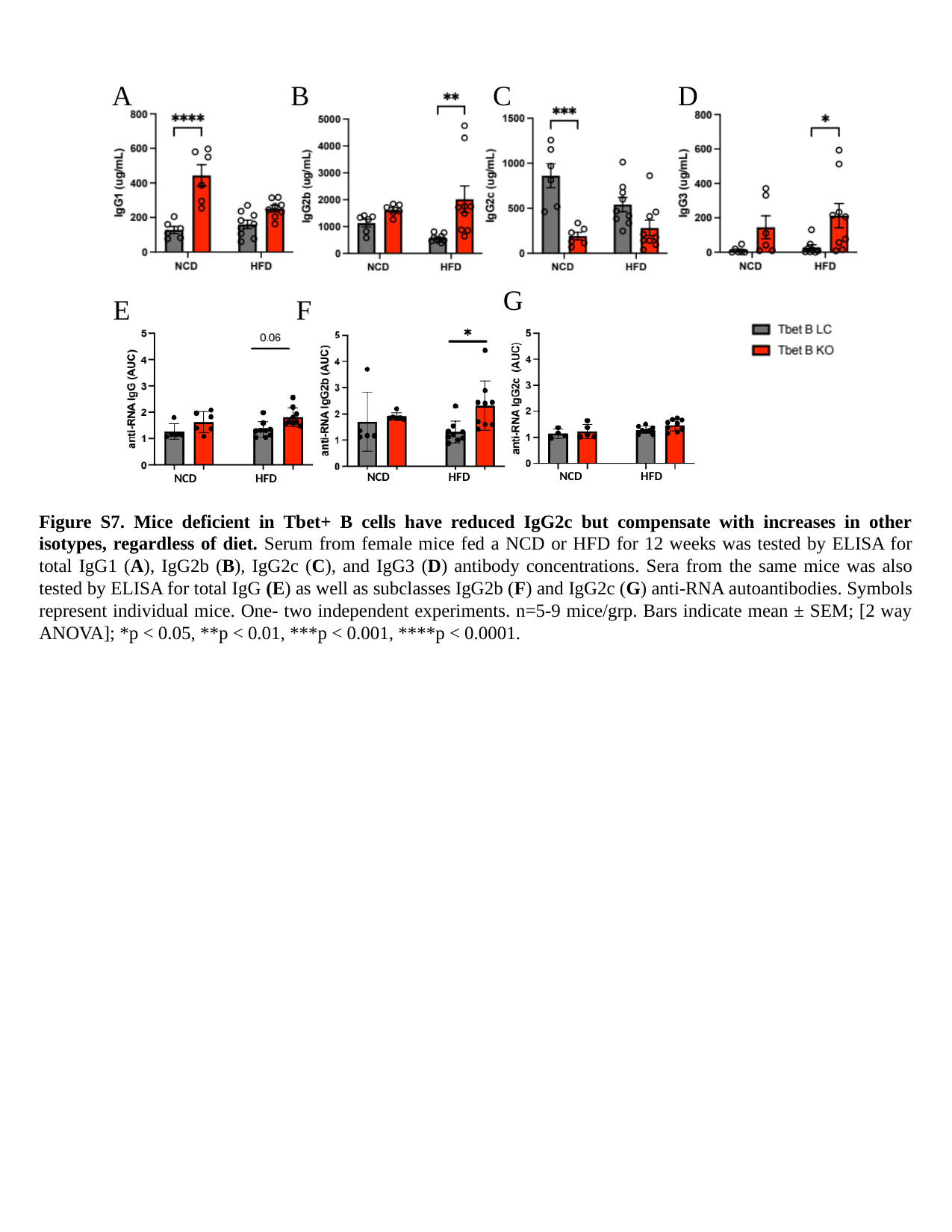

A
B
C
D
G
E
F
NCD HFD
NCD HFD
NCD HFD
Figure S7. Mice deficient in Tbet+ B cells have reduced IgG2c but compensate with increases in other isotypes, regardless of diet. Serum from female mice fed a NCD or HFD for 12 weeks was tested by ELISA for total IgG1 (A), IgG2b (B), IgG2c (C), and IgG3 (D) antibody concentrations. Sera from the same mice was also tested by ELISA for total IgG (E) as well as subclasses IgG2b (F) and IgG2c (G) anti-RNA autoantibodies. Symbols represent individual mice. One- two independent experiments. n=5-9 mice/grp. Bars indicate mean ± SEM; [2 way ANOVA]; *p < 0.05, **p < 0.01, ***p < 0.001, ****p < 0.0001.

### Slide 9
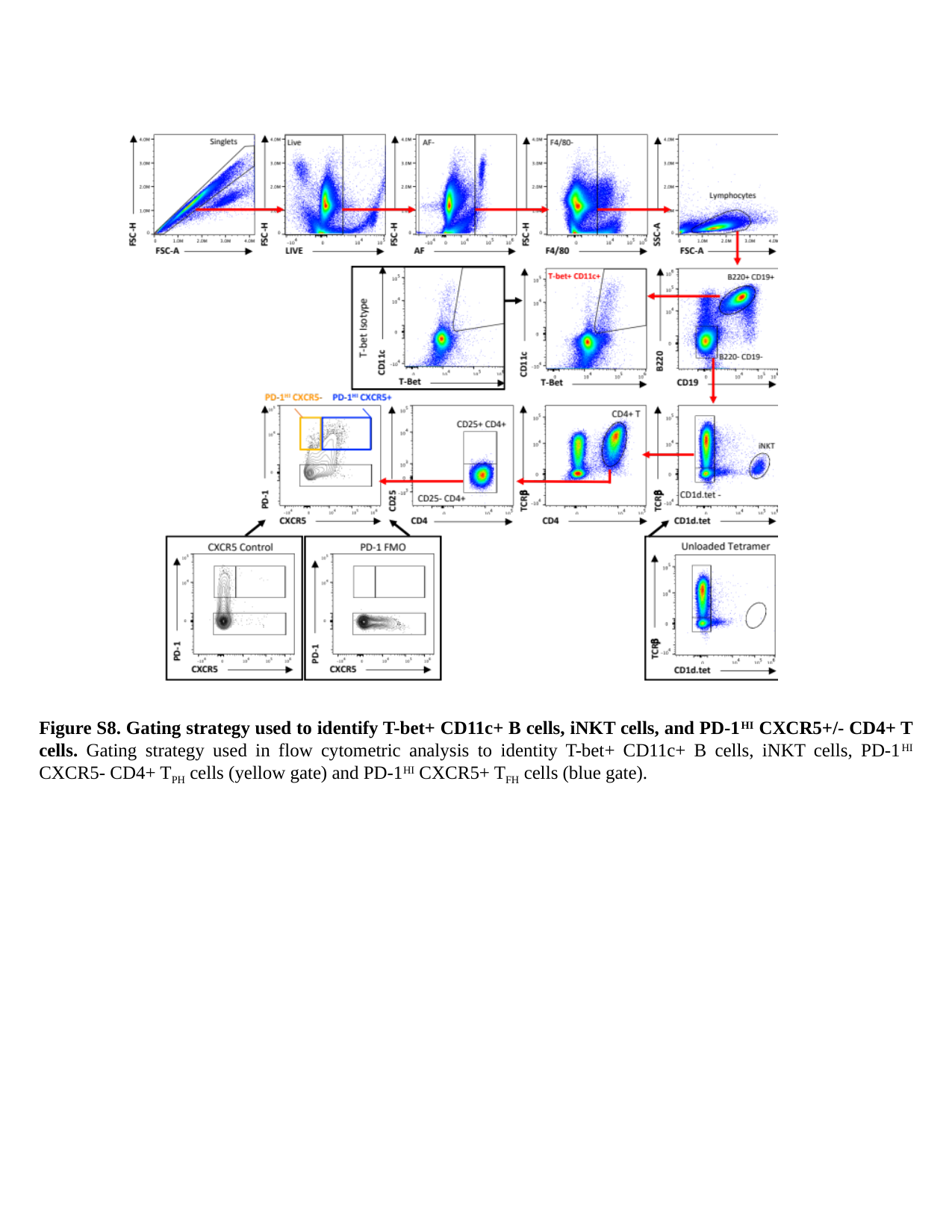

Figure S8. Gating strategy used to identify T-bet+ CD11c+ B cells, iNKT cells, and PD-1^HI^ CXCR5+/- CD4+ T cells. Gating strategy used in flow cytometric analysis to identity T-bet+ CD11c+ B cells, iNKT cells, PD-1^HI^ CXCR5- CD4+ T_PH_ cells (yellow gate) and PD-1^HI^ CXCR5+ T_FH_ cells (blue gate).
